# Individual bacterial taxa drive colonisation resistance to methicillin-resistant *Staphylococcus aureus* in human nasal microbiome samples

**DOI:** 10.64898/2026.01.07.698121

**Authors:** Laura Brülisauer, Mathilde Boumasmoud, Anaëlle Fait, Katia R. Pfrunder-Cardozo, Silvio D. Brugger, Alex R. Hall

**Affiliations:** Department of Environmental Systems Science, Institute of Integrative Biology, ETH Zurich, Zurich, Switzerland; Department of Infectious Diseases and Hospital Epidemiology, University Hospital Zurich, University of Zurich, Zurich, Switzerland

## Abstract

Identifying bacterial interactions that determine susceptibility of human microbiomes to colonisation by pathogenic bacteria has important implications for understanding health and disease and, consequently, improving treatment and prevention. Here, we show how microbiome composition of healthy-human nasal passage samples determines susceptibility to colonisation by methicillin-resistant *Staphylococcus aureus* (MRSA) using a replicated microcosm approach. We find that variable MRSA population growth among samples from different individuals is associated with differences in microbial community composition. To identify individual taxa contributing to in vitro colonisation resistance, we isolate bacteria from inhibitory samples and measure their effects on MRSA in co-culture. This reveals strong MRSA inhibition by multiple Enterobacteriaceae isolates, including some which fully suppress MRSA growth in our system. Finally, by assembling nasal model communities reflecting the composition of natural nasal microbiome samples and combining drop-in and drop-out designs, we show that individual taxa can drive community-level resistance to MRSA growth. Together, our results demonstrate causal links between taxonomic composition of nasal microbiome samples and resistance to *S. aureus*, informing potential new microbiome-targeted interventions to manage *S. aureus* colonisation.

## Introduction

The human nasal passages host a microbial ecosystem that serves as primary reservoir for *Staphylococcus aureus*, and influences respiratory health and disease [1,2]. *S. aureus* frequently colonises the nasal passages and is sometimes referred to as pathobiont [3,4], because it can shift from commensal behaviour to causing infections [2,5–8]. Up to 30% of the human population are persistently colonised by *S. aureus,* placing them at higher risk of subsequent infection by the nasal strain [2,6–12]. Nasal carriage of methicillin-resistant *S. aureus* (MRSA), in particular, can lead to infections with limited treatment options and act as a source of MRSA dispersal [13–15]. Increasing evidence links susceptibility to *S. aureus* colonisation with nasal microbiome composition, suggesting resident nasal microbiota play an important role in conferring colonisation resistance to pathobionts [3,7,16–19]. However, identifying causal links between presence/absence of individual bacterial taxa in microbial communities and susceptibility to nasal *S. aureus* or MRSA colonisation remains a major challenge. Addressing this gap is necessary to improve our fundamental understanding of person-to-person variation in nasal microbiome composition and could inform development of microbiome-based interventions. Such alternatives to conventional antimicrobial-based approaches for managing *S. aureus*/MRSA colonisation are especially relevant in light of the global antimicrobial resistance crisis [20–22].

Current knowledge about bacterial interactions in the nasal microbiome and their relevance for *S. aureus* colonisation is largely based on observational studies and pairwise in vitro experiments. For example, negative correlations between nasal *S. aureus* carriage and the presence of certain commensal species have been reported [3,17,19,23–31]. Such analyses provide important information, but do not identify causal relationships between the presence of individual taxa and susceptibility to colonisation, which would require replicated and controlled experiments with human microbiomes. On the other hand, previous work cultivating relevant strains in vitro revealed direct *S. aureus* inhibition by nasal commensals and investigated underlying molecular mechanisms [24,25,28,32–41]. Such advances strengthened the evidence that microbiota composition contributes to colonisation susceptibility. However, the direct relationship between composition of multispecies microbial communities associated with human nasal passages and susceptibility to colonisation by incoming *S. aureus* or MRSA remains incompletely understood. This question of translation from pairwise cultivation to communities of more than two species is important, because community-level effects often differ from the simple sum of pairwise interactions [42–44]. This likely also applies to the nasal microbiome [33]. To address this gap, we establish an experimental framework linking natural variation in human nasal-microbiome composition with susceptibility to MRSA growth. Eventually, this type of information could improve our understanding of which taxa contribute to individual-level colonisation risk and inform microbiota-oriented treatments [45].

We used a replicated microcosm system to measure colonisation resistance to MRSA in nasal microbiome samples from healthy humans. We interpret colonisation resistance [46] in our system as the ability of a microbiome sample to inhibit growth of an introduced MRSA strain. This allowed us to test three main hypotheses. First, we hypothesized microbiome samples that already contain other, resident strains of *S. aureus* would show relatively strong inhibition of MRSA. This hypothesis is motivated by the ecological principle of competitive exclusion [47] and earlier observations that most *S. aureus* carriers are colonised by a single strain [48]. Second, we hypothesized variable colonisation resistance to MRSA among samples from different individuals is linked with differences in resident microbial community composition (aside from the presence/absence of resident *S. aureus*). Our rationale here is based on past observations that other species inhibit *S. aureus* in pairwise cultivation [24,25,30,32,40], and that nasal microbiome composition differs between *S. aureus* carriers and non-carriers [3,16–19,23–25]. We then assessed the contribution of individual taxa to community-level colonisation resistance, by isolating bacterial strains from inhibitory communities and quantifying their effects on MRSA growth in reconstructed model communities [49]. This allowed us to test a third hypothesis: that individual isolates conferring strong inhibition of MRSA in pairwise cultivation would contribute to colonisation resistance in communities representative of natural human nasal-passage microbiomes.

## Materials and methods

### Sampling human nasal microbiomes

We recruited healthy volunteers aged 18 years or older who had not taken antibiotics in the past three months or nasal spray in the past six weeks, not tested positive for SARS-CoV-2 in the past two months, and not undergone surgery in the past six months. After giving written consent, volunteers were instructed to collect two consecutive nasal swabs of the lower third of the anterior nares (1-2 cm depth) on each side under our supervision using a commercial sampling kit (eSwab 490CE.A, Copan, Italy). We vortexed for 2 min at maximum speed, then pooled the four swabs from each donor, resulting in one sample per donor. Samples were fully anonymised and randomised. This study was confirmed as outside the scope of the Human Research Act and therefore exempt from authorisation (Cantonal Ethics Commission Zurich; BASEC-Nr.: Req-2023-OO678).

### Sample pre-screening and processing

To pre-screen samples for MRSA and resident *S. aureus* (MSSA), we plated 100 µL nasal swab sample suspended in Transport Medium on both CHROMagar™ MRSA (MRSA agar) and CHROMagar™ Staph aureus (SA agar) (CHROMagar^TM^, France) and incubated overnight, while keeping suspended samples and sterilised aliquots at 4 °C. According to pre-screening results, we excluded samples that tested MRSA-positive and randomly selected 12 samples, including six that screened positive for resident MSSA and six that screened negative for resident MSSA. For each of the 12 samples, we stored one 500 µL aliquot with 10% glycerol for bacterial isolation and one 500 µL aliquot without glycerol for sequencing at −80 °C. We made a sterilised version of each sample by autoclaving an aliquot (15 min, 121 °C).

### Inoculation of nasal microcosms and bacterial enumeration

We seeded 230 µL each live / sterile sample (each pooled nasal swab sample split into 3 replicates per sample and condition) into a 5 mL tube (Falcon) containing 890 µL basal medium (RPMI medium 1640 without phenol red, Fisher Scientific, supplemented with 1% brain-heart infusion broth, BHI, Sigma-Aldrich). For a pre-conditioning step, we incubated the resulting 72 microcosms (plus negative controls: only basal medium with sterile Amies Transport Medium) 2 h at 37 °C shaking (180 rpm). Then, we inoculated microcosms with the focal MRSA strain JE2 [50], adding 10 µL of independent overnight cultures in basal medium, diluted 1:10’000 in phosphate-buffered saline (PBS), resulting in a mean inoculum of 413.8 ± 312.4 CFU. We aliquoted 130 µL of each microcosm directly after inoculation with focal MRSA (referred to as timepoint 0 h) for bacterial enumeration (see below), resulting a final microcosm volume of 1 mL, which we incubated 48 h (37 °C, aerobic, shaking). For enumeration of MRSA and total *S. aureus,* we spot-plated (10 µl droplets) serial dilutions at 0 h and 48 h on MRSA and SA agar and incubated 24 h (37 °C, aerobic). Additionally, we plated 50 µL from each microcosm on non-selective Columbia agar with 5% sheep blood (COS, Fisher Scientific) using glass beads (undiluted and 10^−1^ dilution at 0 h, 10^−5^ and 10^−6^ dilutions at 48 h). For each microcosm, we stored one 300 µL aliquot with 20% glycerol for isolation of bacteria and one 500 µL aliquot without glycerol for sequencing.

### 16S rRNA gene sequencing

To assess community composition after sample collection and after the microcosm experiment, we stored 500 µL aliquots of the 12 nasal swab samples and 500 µL aliquots of each post-incubation microcosm at −80 °C. DNA extraction was performed by Zymo Research Europe using the ZymoBIOMICS®-96 MagBead DNA Kit (Zymo Research, USA). We obtained low DNA yield (< 20 ng) for three nasal swab samples (D02, D03 and D12), likely due to low input biomass in the sample, and therefore excluded these samples from further analyses. For all remaining samples (nasal swab samples and final microcosm samples), full-length amplification of the 16S rRNA gene (V1-V9) using primers 27F (5’-AGA GTT TGA TCC TGG CTC AG-3’) and 1492R (5’-CGG TTA CCT TGT TAC GAC TT-3’), library preparation and long-read amplicon sequencing with the GridION instrument (Oxford Nanopore Technologies, MinION R10.4.1 flow cell) were performed by Eurofins Genomics (Germany). We included the Microbial Community Standard (D6300) and the Microbial Community DNA Standard (D6305) from ZymoBIOMICS to assess potential extraction and sequencing biases.

We obtained a higher read count (674’378 ± 168’537, mean ± s.d.) for final microcosm samples compared to unprocessed nasal swab samples (243’105 ± 188’146), presumably because the latter contained lower biomass. To facilitate downstream processing, we generated a random subset of each sample containing 100’000 reads using Seqtk v1.4-r122 (https://github.com/lh3/seqtk). We inspected quality profiles with FastQC v0.12.1 [51] and filtered reads with Filtlong v0.2.1 (https://github.com/rrwick/Filtlong) using the following parameters: length > 1000 bp and < 1600 bp, mean quality score > 20. We used the Emu pipeline v3.4.5 [52,53] with the Emu default database [54–56] for taxonomic abundance tables for each sample and analysed them using phyloseq v1.53.0 [57] in RStudio (R v4.5.1 [58]). For analysis, we retained all taxa with relative abundance >0.2% in at least one sample. This threshold was based on the Microbial Community Standard, which indicated taxa with relative abundances <0.2% are likely spurious.

### Isolation of bacteria from nasal swabs

We isolated examples of each visually distinguishable colony morphotype from each sample plated on COS agar (incubated 48 h, 37 °C, aerobic) and SA agar (incubated 24 h, 37 °C, aerobic) at the start of the experiment. For samples with high Enterobacteriaceae prevalence in the 16S rRNA gene sequencing data, we plated frozen aliquots of the respective timepoint (0 h or 48 h) on chromatic Mueller-Hinton agar (MH agar) (Liofilchem, Italy) and isolated morphologically different colonies after 48 h incubation (37 °C, aerobic). We identified species from colonies using MALDI-TOF. Briefly, we deposited single colonies on a steel plate and treated them with 70% formic acid before analysis in a Bruker Microflex MS system (Bruker Daltonics, Germany) with the alpha-cyano-4-hydroxycinnamic acid matrix. Species assignment was considered correct if the score was ≥2 in two independent identifications. We stored all isolates either (a) belonging to different species or (b), belonging to the same species, but exhibiting a different morphotype, at −80 °C by suspending colonies in PBS with 40% glycerol.

### Pairwise inhibition assays

To test for inhibition of MRSA strain JE2 in co-cultures with isolates from samples associated with MRSA-colonisation resistance, we followed a slightly adapted protocol as for the microcosm experiment above. We inoculated 10 µL of 1:100-diluted overnight culture (or basal medium, for the MRSA pure culture control) in PBS for each isolate (three independent overnight cultures per isolate) into 1120 µL basal medium combined with Amies Transport Medium (4:1 ratio). After 2 h pre-conditioning (37 °C, aerobic, shaking), we added 10 µL of 1:10’000-diluted overnight culture of focal MRSA in PBS (independent cultures for each replicated microcosm), and aliquoted 140 µL for plating timepoint 0 h, resulting in a final volume of 1 mL per microcosm. We incubated all microcosms (48 h, 37 °C, aerobic, shaking), then plated dilutions on MRSA agar and either SA agar for microcosms containing resident *S. aureus* isolates or MH agar for microcosms containing Enterobacteriaceae.

### Membrane-separated co-culture assay

To test whether physical contact with *C. koseri* is necessary for MRSA inhibition, we used a customisable 96-well plate (Versarys, Switzerland) with pairs of wells connected directly or separated by membranes (0.2 µm pore size) that allow exchange of medium. We inoculated individual wells containing 180 µL fresh basal medium with 20 µL diluted overnight cultures of MRSA or *C. koseri* at starting densities between 10^6^-10^7^ CFU/mL (higher than in previous experiments to increase sensitivity for detecting negative population growth) and incubated (48 h at 37°C). We measured bacterial abundance by sampling 10 µL per well at 0, 24, 48 h and plating dilutions on MRSA agar and MH agar.

### Synthetic model communities

Based on the full-length 16S rRNA gene sequencing data from nasal swab samples, we created four model communities with a composition representative for nine donor samples (excluding D02, D03 and D12 samples due to insufficient DNA yield; see Fig. 2a). To compile each of the four model communities (Fig. S8), we selected all isolates present in our collection that:

a. were isolated from plates of nasal microcosms at the start of the experiment and detectable in the 16S rRNA gene sequencing data of the corresponding fresh nasal swab, or
b. constituted >70% of the final community according to 16S rRNA gene sequencing data in all 3 replicated nasal microcosms and were isolated post-hoc by selective plating.

To avoid inter-individual effects, we only used isolates originating from a single, representative sample for each model community (MC_D01, MC_D04, MC_D07, MC_D10). The target total inoculum of each model community was 10^5^ CFU/mL, with the isolate of the dominating species (according to the 16S rRNA gene sequencing data of the fresh nasal swabs) constituting 70% of total abundance and the sum of other isolates accounting for the remaining 30% (Table S2).

Model communities derived from D04 and D07 both included a resident *C. koseri* strain. To test the contribution of *C. koseri* in these communities, we used a ‘drop-out’ approach: including a treatment group where *C. koseri* was not included in the community, comparing this with the complete community (Fig. 5). Abundance of the other isolates in the drop-out communities remained the same as in the unmanipulated communities. The other two model communities, derived from D01 and D10, did not naturally contain detectable *C. koseri.* In these communities we used a ‘drop-in’ approach, comparing unmanipulated communities to versions supplemented with 10^5^ CFU/mL of the *C. koseri* isolate from D07, keeping the abundances of all other isolates the same. See Table S2 for further details.

### Statistical analysis

We performed statistical analyses using R in RStudio (R v4.5.1 [58]). MRSA and total *S. aureus* abundances were quantified by plating serial dilutions, resulting in a limit of detection (LOD) of 100 CFU/mL, indicated in figures. For quantitative analyses, we substituted values below LOD with LOD/2 (corresponds to 50 CFU/mL) and used log_10_-transformed CFU values. Where required due to non-normally distributed data or unequal variances across groups, we used non-parametric methods or rank-transformation followed by ANOVA to test main effects. For experiments where there was a clear separation into groups with positive growth vs those below LOD, we used logistic regression.

## Results

### Susceptibility to MRSA colonisation varies among samples from different individuals

To measure colonisation success of an incoming MRSA strain in natural nasal communities, we collected nasal-swab samples from healthy human volunteers. We seeded 12 samples into replicated nasal microcosms with basal medium (RPMI 1640 + 1% BHI, see Methods; Fig. 1a), including six samples that contained resident methicillin-susceptible *S. aureus* (MSSA), and six that did not (according to a pre-screening step; see Methods). We then inoculated all microcosms with the focal MRSA strain JE2, derived from epidemic clone USA300 and widely used in experimental studies [50], and measured its abundance after 48 h incubation. Final MRSA abundance was reduced in nasal microcosms seeded with live microbiome samples compared to equivalent microcosms containing sterilised versions of the same samples (*p* < 0.001, Wilcoxon signed-rank test, using median for each sample; Fig. 1b), indicating the presence of a live resident community on average decreases MRSA growth. The degree of colonisation resistance to MRSA in live microcosms, taken here as final MRSA abundance, varied among microcosms seeded with samples from different donors (two-way ANOVA on rank-transformed data: *F*_10,24_ = 14.57, *p* < 0.001; Fig. 1b). Colonisation resistance was generally higher in samples that already contained other, resident strains of *S. aureus* (D07-D12), compared with the resident-*S. aureus*-negative samples (D01-D06) (two-way ANOVA on rank-transformed data: *F*_1,24_ = 41.22, *p* < 0.001). In these resident-*S. aureus*-positive communities, total *S. aureus* abundance consistently exceeded focal MRSA abundance, indicating resident *S. aureus* was still present at the end of the experiment, although in variable abundances depending on the donor sample (Fig. S1a). In some samples, both total *S. aureus* and MRSA showed little or even negative population growth over time (Fig. S1b). These results indicate resident microbiota can fully inhibit population growth of incoming MRSA, but the strength of this effect varies among samples from different individuals.

**Figure 1:**
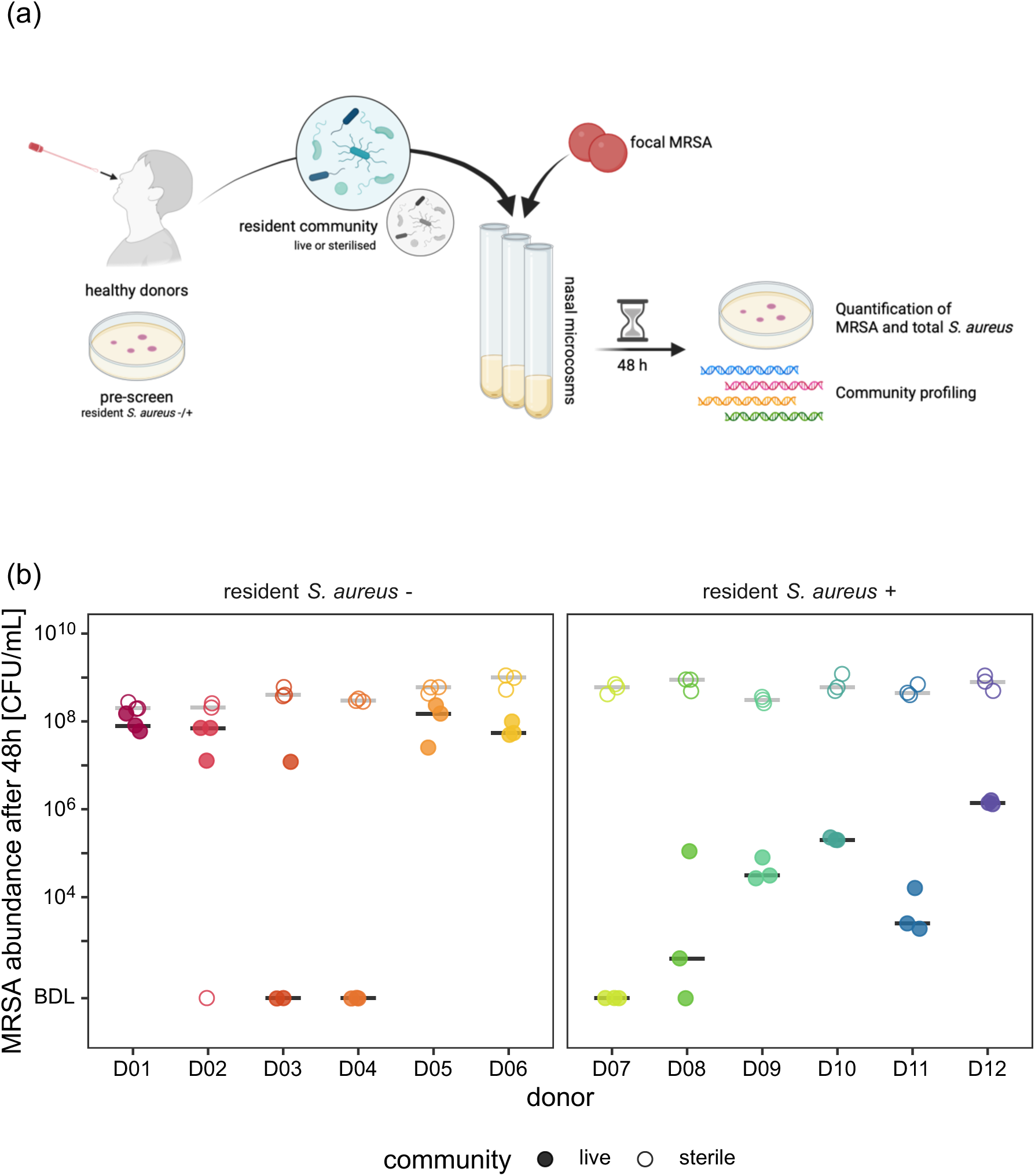
MRSA colonisation success in nasal microcosms varies among microbiome samples from different individuals. (a) Experimental set-up: We collected nasal swab samples from 12 healthy human donors and created replicated nasal microcosms containing either live or sterilised sample. Subsequently, we introduced focal MRSA strain JE2 (derived from the epidemic clone USA300) and measured its abundance after 48 h as a proxy for colonisation resistance of the nasal microbial community in each microcosm. Schematic created with BioRender.com. (b) MRSA abundance after 48 h in microcosms containing live nasal swab samples (filled points, *n* = 3 per donor) and sterilised versions of the same samples (empty points, *n* = 3 per donor). The median across replicated microcosms is shown as a line for each donor (black = live microcosms, grey = sterile microcosms).

### Colonisation resistance is linked with final community composition

We investigated whether observed variability in MRSA colonisation outcomes among samples from different donors was correlated with differences in resident community composition using species-level community profiling by full-length 16S rRNA gene sequencing. Community composition varied markedly among donors (Fig. 2a) and reflected three out of seven previously described community state types [19], suggesting our samples captured a significant proportion of the diversity present in nasal microbiomes among the general population. We found complete MRSA inhibition (final abundance below detection limit) in microcosms seeded with samples from four different donors (Fig. 1b). The final composition of each fully resistant microcosm was dominated by a single Enterobacteriaceae species: *Klebsiella pneumoniae* (D03), *Citrobacter koseri* (D04 and D07), and *Enterobacter hormaechei* (D08) (Fig. 2b). Principal coordinate analysis supported this link between high relative final abundance of Enterobacteriaceae and final MRSA abundance (Fig. 2c). Beyond complete colonisation resistance to incoming MRSA, the two communities among these which included a resident *S. aureus* strain at the start (D07 and D08) showed a substantial decline in total *S. aureus* abundance over time, resulting in lower final abundance compared to other resident-*S. aureus*-positive communities (Fig. S1). This suggests resident communities from D07 and D08 supported little growth of both focal MRSA and resident *S. aureus* in our experiment. In summary, community composition, particularly the presence of various Enterobacteriaceae species, was associated with colonisation resistance to MRSA in nasal microcosms.

**Figure 2:**
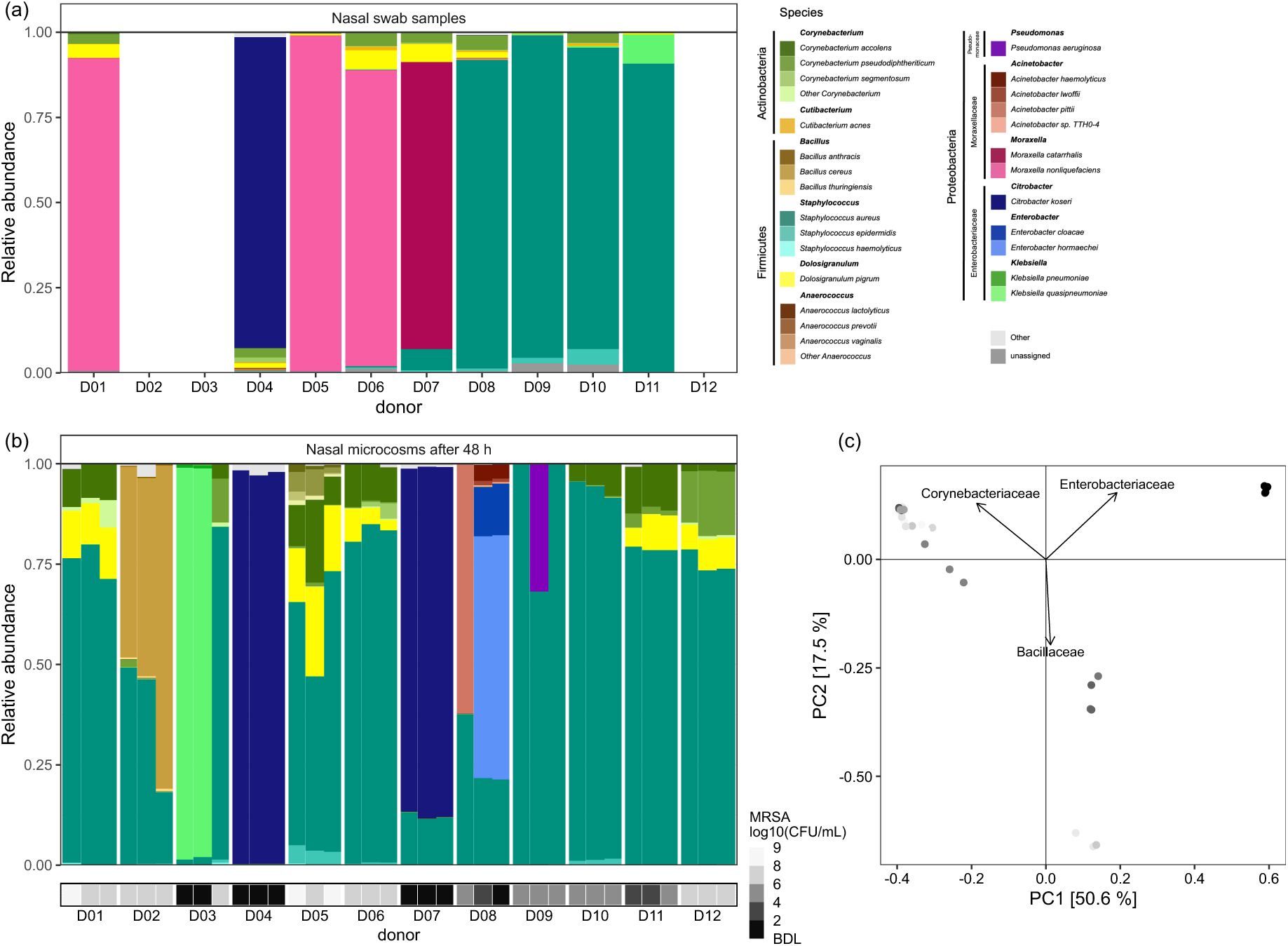
MRSA colonisation resistance is associated with high relative abundance of Enterobacteriaceae in microcosms after 48 h. (a) Species-level community composition of the unmanipulated nasal swab samples after collection assessed with full-length 16S rRNA gene sequencing. Samples from donors D02, D03 and D12 are omitted because of insufficient DNA yield (< 20 ng, leading to < 20’000 reads), which is likely attributed to low biomass commonly found in nasal swab samples. (b) Species-level composition in replicated live microcosms after 48 h (*n =* 3 per sample). For reference, we show MRSA abundance after 48 h in each microcosm, used here as a proxy for colonisation resistance (see Fig 1), as a heatmap. Note that in microcosms, bars for *S. aureus* reflect the sum of focal MRSA and any resident *S. aureus* (present in D07-D12). These cannot be differentiated with 16S rRNA amplicon sequencing, but data separating MRSA from total *S. aureus* by selective plating is provided in Fig. S1. (c) Principal coordinate analysis (PCoA) on family-level Bray-Curtis dissimilarity supports the correlation between Enterobacteriaceae prevalence in resident microbiota and final MRSA abundance. We excluded *S. aureus* from the dataset before merging species at family level to avoid growth of the focal strain affecting the analysis. Each point represents one microcosm, the colour scale indicates MRSA abundance after 48 h (colour gradient same as in (b)). The arrows indicate the contribution of all families with an r^2^ > 0.5 and *p* < 0.01 (determined with the envfit function from vegan package v2.7-1 in R [74]).

### Isolates from inhibitory communities suppress MRSA in pairwise co-culture

To identify which individual taxa contributed to community-level colonisation resistance, we tested MRSA inhibition by individual bacterial isolates in the absence of the community background, using pairwise co-cultivation. Given the higher average colonisation resistance in resident-*S. aureus*-positive communities observed above (Fig. 1b), we first isolated one resident *S. aureus* strain from each of the six positively screened donor samples (D07-D12). We then co-cultured each resident *S. aureus* isolate with the focal MRSA. All resident *S. aureus* (rSau) isolates decreased MRSA growth compared to pure MRSA culture (Fig. 3a). We detected no tendency for stronger inhibition depending on resident-isolate sequence type, *agr* type (Fig. S2, Table S1), average nucleotide identity with JE2 (Fig. S3b), or relative pure-culture growth rate (Fig. S3a).

**Figure 3:**
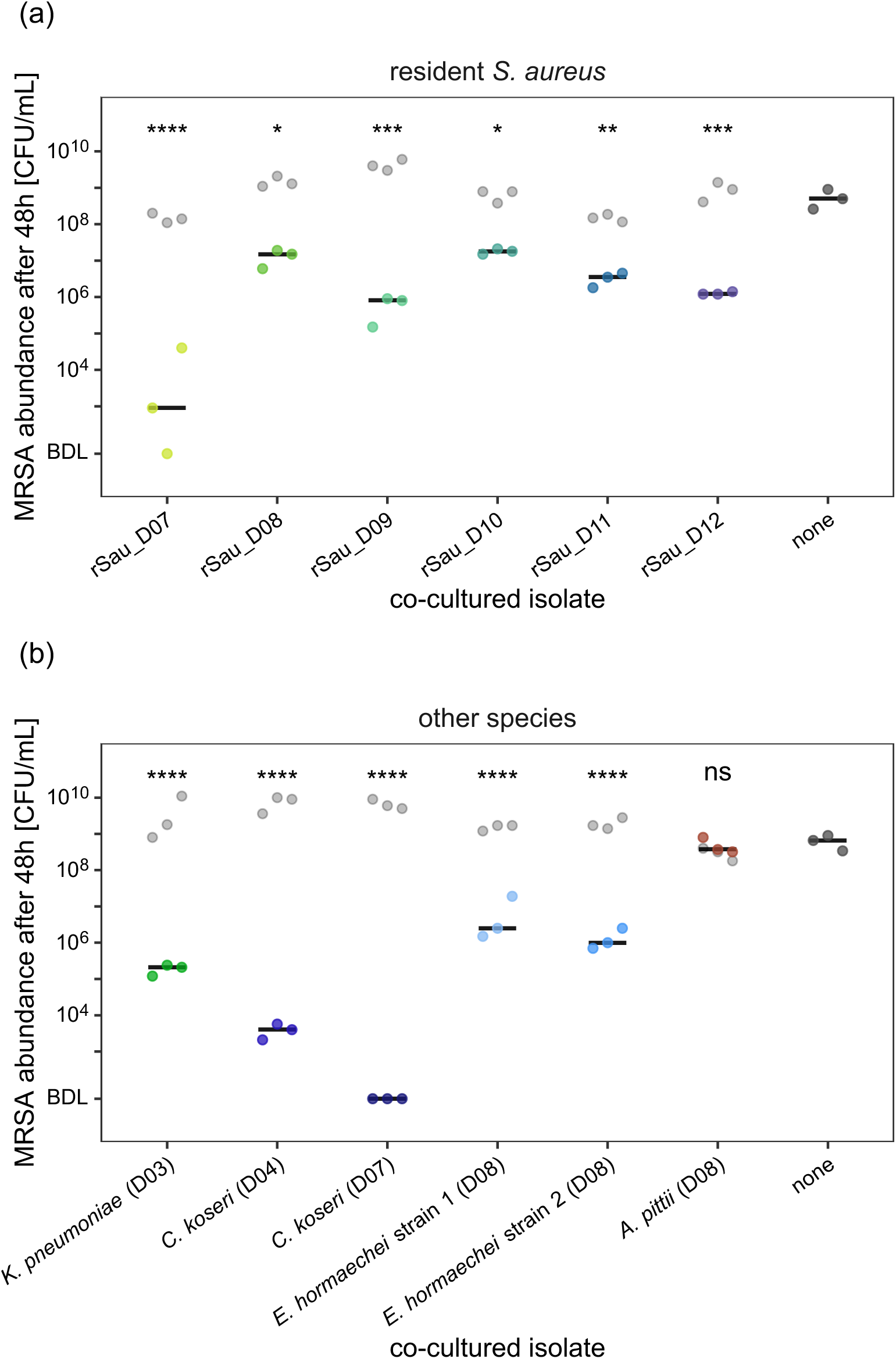
Isolates from inhibitory communities inhibit MRSA growth in pairwise co-culture. Both (a) resident *S. aureus* (rSau) isolates from donors D07-D12 and (b) other species including different Enterobacteriaceae isolates reduce MRSA growth (coloured points) in absence of their native community in liquid co-culture (*n* = 3) compared to pure MRSA culture (dark grey points, labelled as “none”). Co-culture with the non-Enterobacteriaceae species *A. pittii* (*Moraxellaceae*) is included for reference. Light grey points indicate the final abundance of the respective co-cultured isolate (coloured and grey points are grouped by replicate). Black lines indicate the median MRSA final abundance across replicates. Asterisks display significance levels of all unpaired two-sample *t*-tests comparing MRSA (log-transformed abundance) in co-cultures to MRSA pure culture after correction for multiple testing by sequential Bonferroni (*p* < 0.0001 ****, *p* < 0.001 ***, *p* < 0.01 **, *p* < 0.05 *).

The strong MRSA inhibition observed in resident-*S. aureus-*negative samples from D03 and D04 and the reduced total *S. aureus* abundance in *S. aureus*-positive samples from D07 and D08 suggested the presence of other taxa that interfere with *S. aureus* growth. Based on the association between colonisation resistance to MRSA and high final Enterobacteriaceae abundance (Fig. 2), we isolated the dominating Enterobacteriaceae species from relevant samples (1×*K. pneumoniae*, 2×*C. koseri*, 2×*E. hormaechei*) and tested MRSA inhibition in pairwise co-culture. As above for resident *S. aureus*, these isolates inhibited MRSA growth to varying degrees, with the two *C. koseri* isolates from D04 and D07 having the strongest effects (Fig. 3b). For reference, we included the non-Enterobacteriaceae species *Acinetobacter pittii* (*Moraxellaceae*), which reached high relative abundance in one microcosm seeded with the D08 sample (Fig. 2b). This revealed no effect of *A. pittii* on MRSA growth, demonstrating successful growth in a community does not necessarily reflect MRSA-inhibitory properties in pairwise co-culture. Consistent with this, isolate growth did not correlate with MRSA inhibition in pairwise co-cultures (Fig. S4).

A further inhibition assay, where resident isolates and MRSA were inoculated side-by-side on agar, revealed MRSA inhibition in proximity of all Enterobacteriaceae isolates (Fig. S5a). By contrast, resident-*S. aureus-*mediated MRSA inhibition observed in liquid co-culture (Fig. 3a) was not visible in side-by-side spot assays (Fig. S5b), suggesting a different mode of action. Cultivating MRSA directly in sterile-filtered Enterobacteriaceae culture supernatants was consistent with inhibitory effects observed above, in that they did not support detectable MRSA growth (Fig. S6). Supplementing supernatants with 10% fresh basal medium partially restored growth (Fig. S6a-b). Together, these findings show Enterobacteriaceae isolates, particularly *C. koseri,* inhibit MRSA under different conditions, suggesting both direct interference (Fig. S5a) and indirect competition, for example via nutrient depletion (Fig. S6), may play a role.

### *C. koseri* kills MRSA in a contact-dependent manner

We further investigated the conditions required for MRSA-inhibition by *C. koseri* D07, which had exhibited the strongest inhibiting effect in pairwise co-culture (Fig. 3b), by comparing MRSA growth in mixed co-cultures vs membrane-separated co-cultures (0.2 µm pore size). In mixed co-culture with *C. koseri* D07, MRSA abundance declined, resulting in complete eradication after 24 h (Fig. 4). By contrast, when separated from *C. koseri* D07 by a membrane, MRSA reached a high median final abundance, only slightly lower than in the pure-culture control (Fig. 4). This suggests a contact-dependent killing mechanism. Consistent with this, *C. koseri* D07 culture supernatant did not induce negative MRSA population growth in an additional experiment where we monitored bacterial density by plating and counting CFUs (Fig. S7), rather than optical density as in Fig. S6. Thus, *C. koseri* D07 caused a net decline in MRSA population size, but this depended on physical contact between the two competitors.

**Figure 4:**
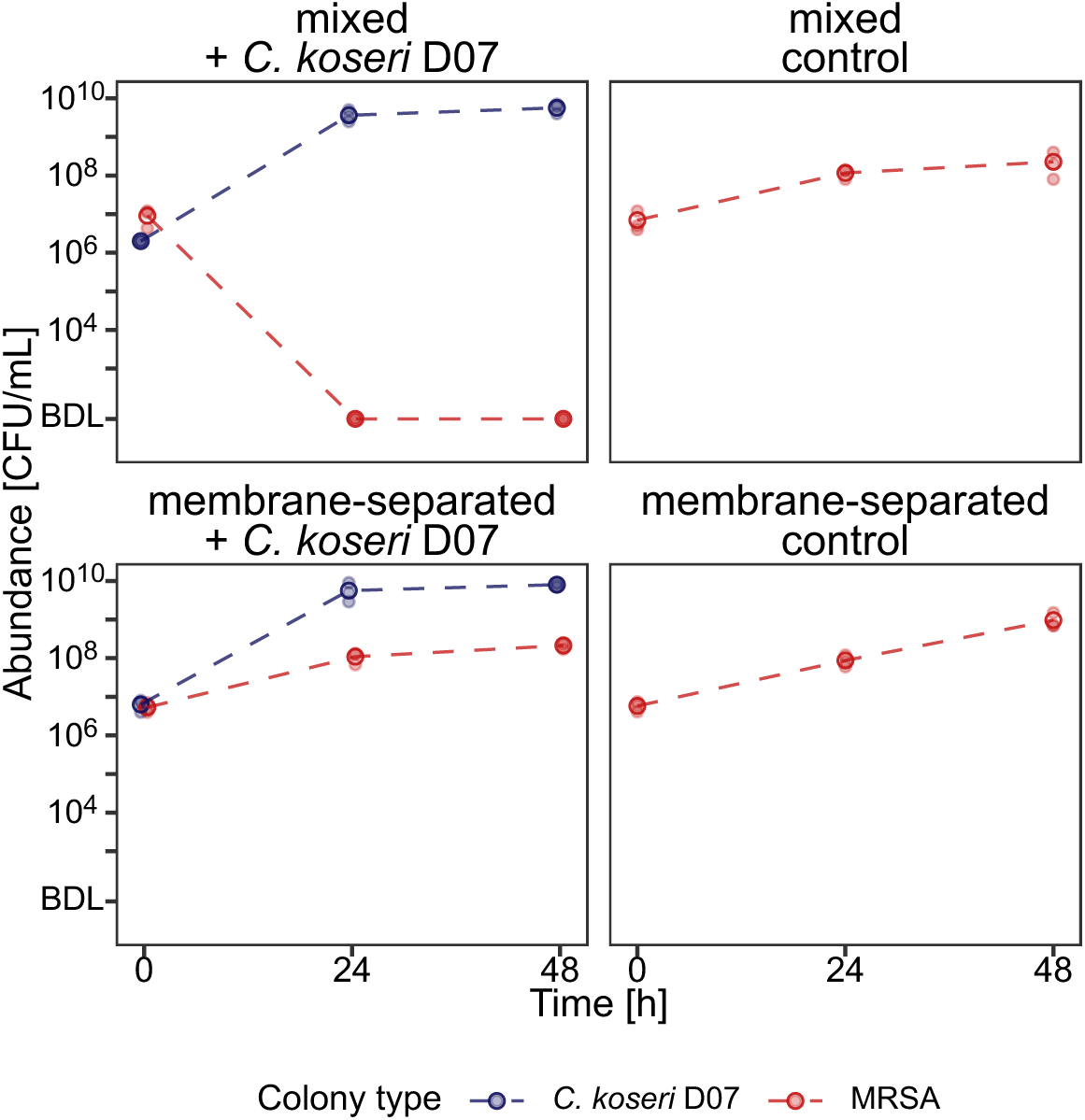
*C. koseri* inhibits MRSA in a contact-dependent manner. MRSA (red) was completely eradicated within 24 h in mixed co-culture with *C. koseri* D07 (blue) (top-left panel). When separated from *C. koseri* D07 by a membrane (0.2 µm pore size), allowing exchange of medium and soluble compounds but not bacterial cells (bottom-left panel), MRSA reached final abundance slightly lower than in the pure culture control (bottom-right panel; median reduction = 0.5 log_10_ CFU; unpaired *t-*test on log-transformed data: *p* < 0.01). Filled points represent individual replicates (*n* = 3), empty circles and dashed lines show median abundance over time.

### Individual taxa determine colonisation resistance in reassembled communities

We next tested whether inhibition of MRSA in pairwise co-culture with *C. koseri* translated to multispecies communities derived from human-microbiome samples. We assembled four nasal model communities (MC_D01, MC_D04, MC_D07, MC_D10), each combining taxa from one of four samples (D01, D04, D07, D10), representative of initial sample composition (Fig. S8, Table S2). The composition of these four samples captured the five most abundant genera across the entire set of samples (*Staphylococcus, Moraxella, Citrobacter, Corynebacterium* and *Dolosigranulum*) and represented three previously described nasal community state types (CSTs) [19]: CST1 (dominated by *S. aureus*; MC_D10), CST2 (dominated by Enterobacteriaceae; MC_D04), and CST6 (dominated by *Moraxella* spp.; MC_D01 and MC_D07).

For the two model communities containing native, resident *C. koseri* (MC_D04 and MC_D07, derived from samples D04 and D07), we compared MRSA growth in presence of the complete community to drop-out versions lacking *C. koseri* (Fig. 5a). MRSA growth in both complete model communities was inhibited to levels comparable to those observed in the respective natural communities (Fig. 1b). Removal of *C. koseri* resulted in much higher MRSA growth (two-way ANOVA on rank-transformed data, effect of *C. koseri*: *F*_1,9_ = 29.45, *p* < 0.001), identifying *C. koseri* as a key driver of colonisation resistance here. For example, the complete MC_D07 (including *C. koseri*) was fully colonisation resistant, but removing *C. koseri* resulted in final MRSA abundance only slightly lower than the pure-culture control. Consistent with microcosms using samples D04 and D07 in our main experiment (Fig. 2b), both unmanipulated model communities (not drop-out versions) were dominated by resident *C. koseri* after incubation (Fig. S9a). Presence of *C. koseri* in model communities was associated with higher final total bacterial abundance (Fig. S9).

**Figure 5:**
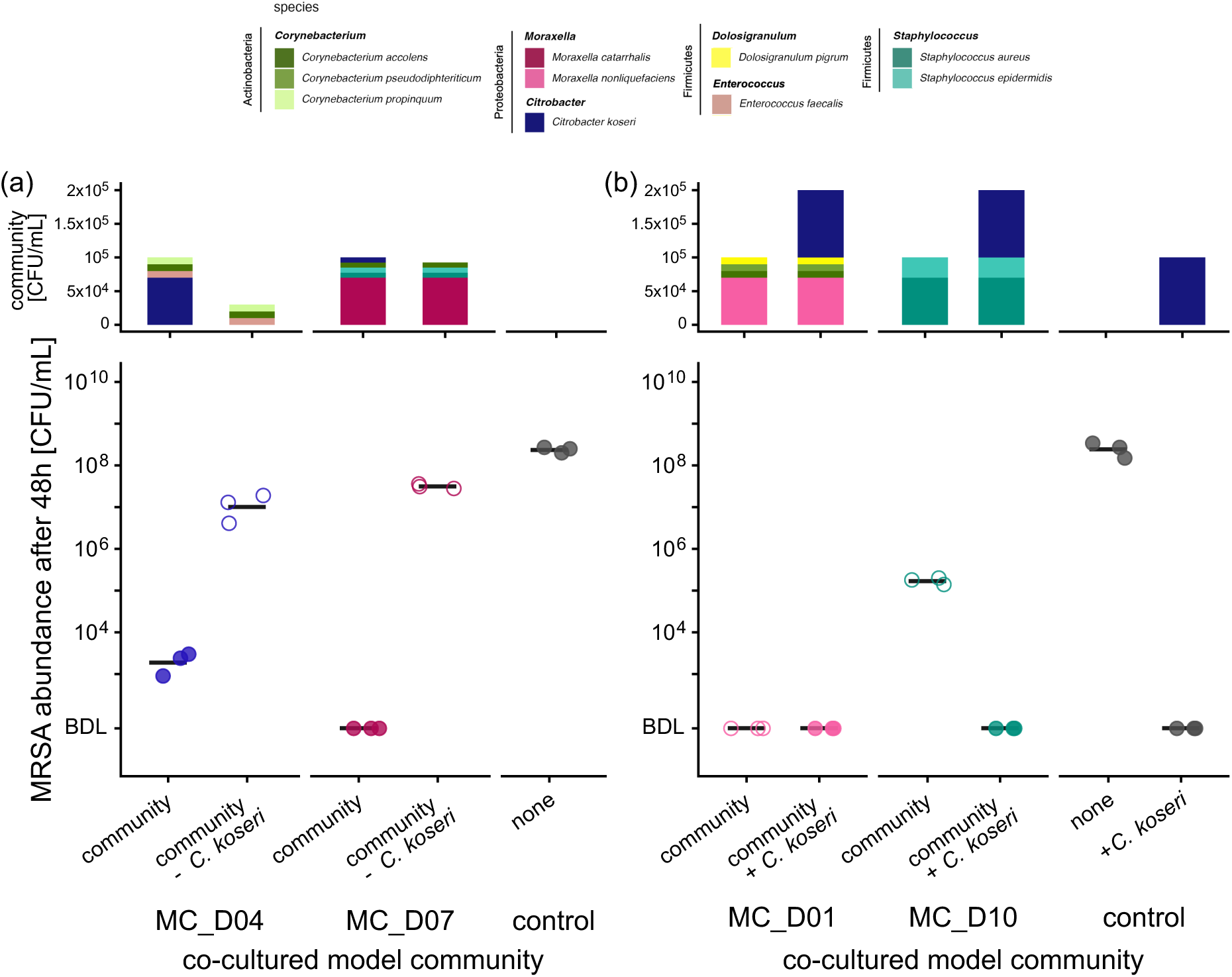
*C. koseri* determines colonisation resistance of assembled communities to incoming MRSA. (a) Final MRSA abundance is decreased when resident *C. koseri* is present, identifying it as a key driver of colonisation resistance to MRSA in the model communities derived from D04 and D07. (b) Adding a *C. koseri* isolate to communities lacking resident Enterobacteriaceae confers colonisation resistance to the otherwise susceptible D10 model community, whereas the D01 model community exhibited colonisation resistance regardless of inoculation with *C. koseri.* Initial community composition is displayed in the top panels (showing the target abundance of each species upon inoculation; see Methods), and final MRSA abundance in the microcosms is shown in the bottom panels.

To test whether introducing *C. koseri* to resident-Enterobacteriaceae-negative communities enhances MRSA-colonisation resistance, we added *C. koseri* D07 to model communities MC_D01 and MC_D10 (Fig. 5b). Both model communities fully inhibited MRSA when inoculated with *C. koseri*. Thus, addition of *C. koseri* decreased the probability of MRSA detection after incubation (logistic regression: log-odds = – 3.56 ± 1.68, *p* = 0.034). Susceptibility of MC_D10 to MRSA colonisation without *C. koseri* agrees with the main microcosm experiment, but the lack of MRSA growth in MC_D01 even without *C. koseri* contrasts the high susceptibility to MRSA colonisation in the D01 nasal-microbiome sample (Fig. 1b). The microcosms seeded with MC_D01 were dominated by *Corynebacterium pseudodiphtheriticum* after 48 h (Fig. S9b). Further experiments revealed *C. pseudodiphtheriticum* to be the key driver of colonisation resistance in MC_D01 (Fig. S10), and that its effects can be influenced by its inoculum density and presence of other species, potentially explaining the divergent outcomes in MC_D01 (Fig. 5b) compared with the corresponding D01 nasal-microbiome sample (Fig. 1b).

To test our main findings under nutrient conditions more representative of nasal secretions, we repeated drop-in/drop-out experiments in synthetic nasal medium (SNM3) [59]. Consistent with results in basal medium (RPMI 1640 + 1% BHI) (Fig. 5), *C. koseri* drove community-level colonisation resistance to MRSA, while the model communities lacking *C. koseri* were even more susceptible to MRSA colonisation than in basal medium (Fig. S11), which was also reflected by final community composition (Fig. S12). In SNM3, susceptibility of MC_D01 without *C. koseri* to MRSA colonisation (Fig. S11b) was in line with the main microcosm experiment (Fig. 1b). In summary, individual taxa, like *C. koseri* or *C. pseudodiphtheriticum,* can determine community-level colonisation resistance to MRSA in vitro.

## Discussion

We established an experimental microcosm approach to test for a causal relationship between nasal-microbiome composition and MRSA growth in human samples, then used it to quantify the contribution of individual taxa to community-level colonisation resistance. We found nasal microbiome samples from healthy humans vary in their resistance to MRSA, and this inter-individual variation is linked with community composition. Pairwise assays and targeted manipulation of model communities demonstrated both intra- and interspecies interactions contribute to MRSA inhibition, and community-level outcomes can be driven by individual taxa.

The starting conditions in microcosms incorporated biotic and abiotic components of nasal-microbiome samples. However, microcosms deviate from the environment in the anterior nares in important ways, such as nutrient composition, host factors, spatial structure and temperature or oxygen gradients. An important limitation is therefore that changes in community composition upon cultivation may reduce the relevance of observed interactions, for example if the conditions favour fast-growing bacteria (e.g., *S. aureus* or Enterobacteriaceae) over other important, but more fastidious nasal taxa. We observed three main types of compositional changes, depending on which sample microcosms were seeded with: 1) CST6 (*Moraxella*-dominated) to CST1 (*S. aureus*-dominated) for samples D01, D05 and D06, attributable to growth of incoming focal MRSA, 2) CST1 (*S. aureus*-dominated) to CST2 (Enterobacteriaceae-dominated) for sample D08, and 3) CST6 (*Moraxella*-dominated) to CST2 (Enterobacteriaceae-dominated) in D07 (Fig. 2a,b). Despite these changes, final community compositions reflect previously described nasal community state types [19,60]. Furthermore, in four samples, the dominating species remained stable during the experiment (*C. koseri* in D04; *S. aureus* in D09-D11), and in other microcosms the relative abundance of other common nasal commensals was stable or increased (e.g., *Corynebacterium* spp. and *D. pigrum* in samples D01, D05 and D06; Fig. 2). Our finding that the fastidious *C. pseudodiphtheriticum* (Fig. S10), not only *C. koseri* (Fig. 5), can drive MRSA inhibition supports the system as capturing some relevant interactions beyond those involving fast-growing Enterobacteriaceae. Furthermore, the robustness of *C. koseri*-mediated inhibition across experimental conditions, including on agar (Fig. S5) and in communities in synthetic nasal medium (Fig. S11), suggests our main findings are not specific to one set of conditions.

Further indication some of our key results are relevant beyond our system comes from past work showing Enterobacteriaceae can be potent *S. aureus-*antagonists. For example, some of the same species were implicated by past work investigating links to *S. aureus-*carrier status with independent cohorts of healthy donors, and using different methodologies to test for in vitro *S. aureus* inhibition, including a nasal epithelium model [24,25]. Similarly, *C. pseudodiphtheriticum*-mediated *S. aureus* inhibition (Fig. S10) is consistent with past work [61]. Potential relevance of these interactions is also supported by evidence Enterobacteriaceae can colonise the nasal passages; indeed, CST2 is dominated by Enterobacteriaceae and negatively associated with *S. aureus* carriage, consistent with our observations [19,60]. Nevertheless, it is important to note CST2 is rare in the human population and typically characterised by high total bacterial abundance and low stability [19,60]. Therefore, any impact of Enterobacteriaceae-*S. aureus* interactions is likely to vary among individuals and potentially over time as well.

We found a nasal *C. koseri* isolate can kill MRSA in a contact-dependent manner. A common molecular mechanism underlying contact-dependent bacterial interactions is type VI secretion system (T6SS) [62–64]. Consistent with this, we detected a putative T6SS^i^ in the isolate’s genome (Table S3). This type of inhibition does not preclude other mechanisms, such as nutrient depletion or other changes to the abiotic environment. Indeed, our supernatant experiments suggest prior Enterobacteriaceae growth changes the abiotic conditions, decreasing *S. aureus* growth (Fig. S6). Although final MRSA abundance was also slightly lower in membrane-separated real-time co-culture with *C. koseri* compared with a pure culture control, MRSA was completely eradicated in mixed co-culture (Fig. 4). This suggests the *C. koseri-*mediated drop in MRSA abundance is mainly driven by contact-dependent inhibition, but other interactions can also contribute. That multiple mechanisms can be involved in Enterobacteriaceae-mediated inhibition is in line with previous work implicating, for example, production of antimicrobials or siderophore-based interactions [24,25,40].

In terms of basic microbial ecology, our results show compositional variation among individual healthy microbiomes, widely documented for the nasal passage and other body sites [19,26,27,29,65–69], causes functional differences in terms of susceptibility to colonisation by incoming bacteria. That multiple taxa, from samples from different individuals, each inhibited MRSA suggests drivers of colonisation resistance vary among individuals. Understanding why some microbiomes resist colonisation, while others remain susceptible, is also relevant for development of microbiome-targeted interventions, for example personalised approaches to decolonisation of patients in at-risk groups or prior to surgery [9,70,71]. Information about causal connections between community composition and susceptibility to *S. aureus* / MRSA colonisation could enable predicting and monitoring impacts of probiotic interventions and assessing recolonisation risks, based on community-composition data for individual patients [21]. Our finding that a single taxon can determine community-level resistance to MRSA is encouraging in this context, in that the presence of a single taxon is likely to be relatively amenable to targeted manipulation, compared with an alternative scenario where colonisation resistance would depend on a particular multi-species composition or higher-order interactions. However, harnessing Enterobacteriaceae-mediated *S. aureus* inhibition for new decolonisation strategies raises important practical considerations, as many Enterobacteriaceae species have pathogenic potential. *C. koseri*, despite being isolated from healthy human microbiomes here and in previous studies [24,40], can cause severe infections [72,73]. Molecular characterisation of inhibitory mechanisms could inform strategies for risk mitigation, for example screening species with similar *S. aureus*-inhibitory characteristics but lower infection risk, isolating inhibitory compounds, or generating attenuated or avirulent strains that retain relevant properties.

## Acknowledgements

We thank the participants of the nasal sampling, Tomas Demeter for advice on bioinformatics, and members of the Pathogen Ecology group and the SDB Lab for feedback and support. This work was funded by the SNSF grant 320030-231258 to Alex R. Hall and the SNSF grant 211422 to Silvio D. Brugger.

## Author contributions

L.B., S.D.B. and A.R.H. conceptualised the project and designed the experiments. L.B., M.B., A.F. and K. R. P. developed the methodology. L.B. collected the samples. L.B. and M.B. performed the experiments. L.B. undertook bioinformatic and statistical analyses and visualised the data. A.R.H. and S.D.B. secured funding. L.B. and A.R.H. drafted the original manuscript, and all authors read, reviewed and approved the manuscript.

## Data and code availability

The 16S rRNA amplicon sequencing data from the twelve nasal microbiome samples, and the genomes of the six resident *S. aureus* isolates and the five Enterobacteriaceae isolates, are uploaded to ENA (European Nucleotide Archive) and will be made accessible upon publication (accession number: PRJEB106874). All other data supporting the findings of this study and the code used for analysis will be available on an open-source repository upon publication.

