## Supplementary Material for "Individual bacterial taxa drive colonisation resistance to methicillin-resistant *Staphylococcus aureus* in human nasal microbiome samples"

### Supplementary Figures

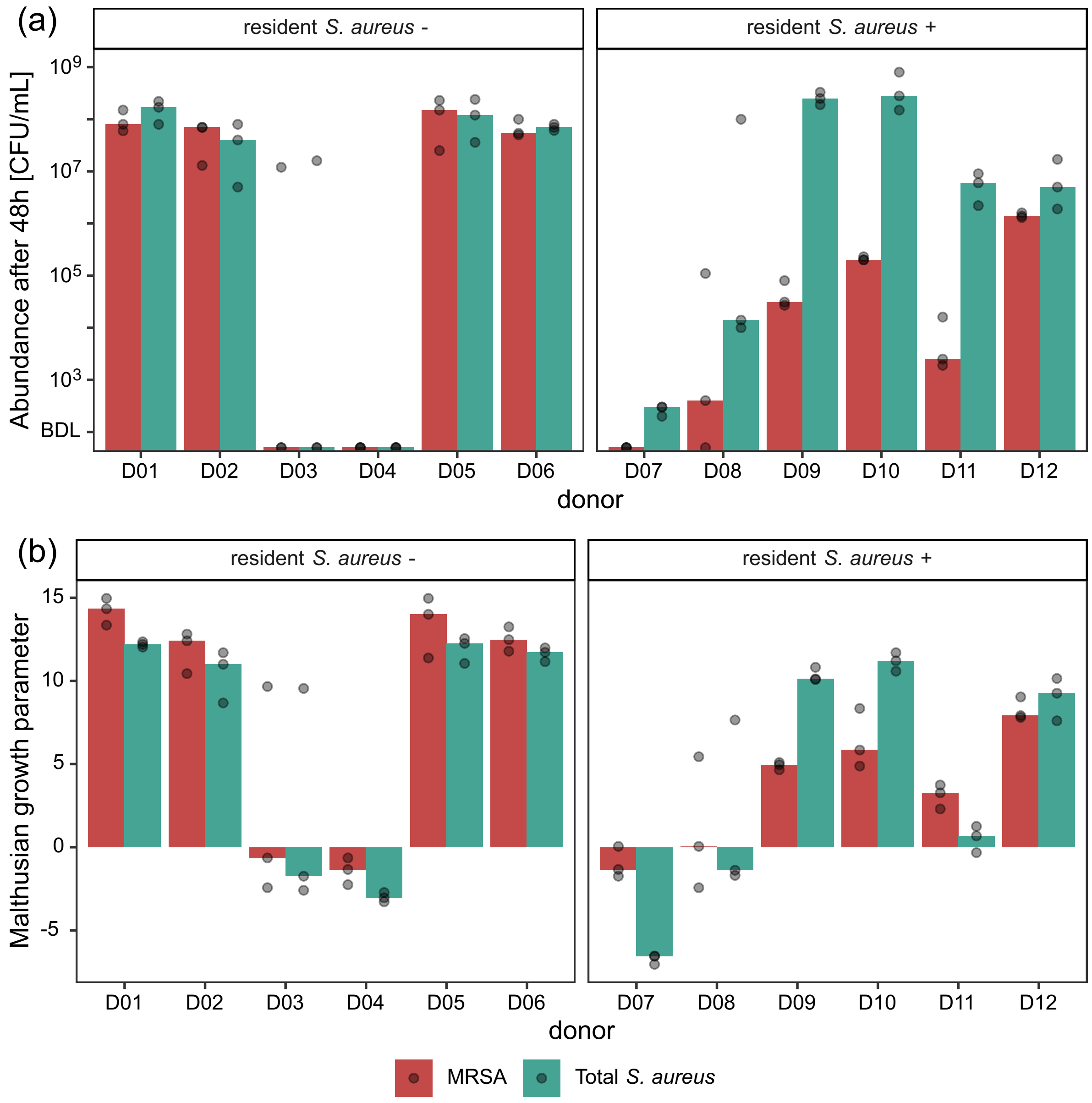


**Supplementary Figure 1: (a) Final abundance of MRSA** (red) **and total *S. aureus*** (turquoise) after 48 h in microcosms from each donor ($n = 3$) **and (b) corresponding Malthusian growth parameter** ($\ln(\frac{\text{final abundance}}{\text{inoculum}})$) [1]. We assessed final MRSA and total *S. aureus* abundance by plating the same dilutions on both MRSA- and *S. aureus*-selective chromatic agar. In resident-*S. aureus*-negative communities (left panels), final abundance of

MRSA and total *S. aureus* are equivalent. In resident-*S. aureus*-positive communities (right panels), final total *S. aureus* abundance depended on the donor (one-way ANOVA on ranktransformed data: $F_{11,24} = 12.69$, $p < 0.001$) and exceeded final MRSA abundance overall ($p = 0.031$, Wilcoxon signed-rank test on medians of replicated microcosms for each donor). The individual replicates are represented by points, bars indicate the median. The detection limit of 100 CFU/mL is indicated as a black line.

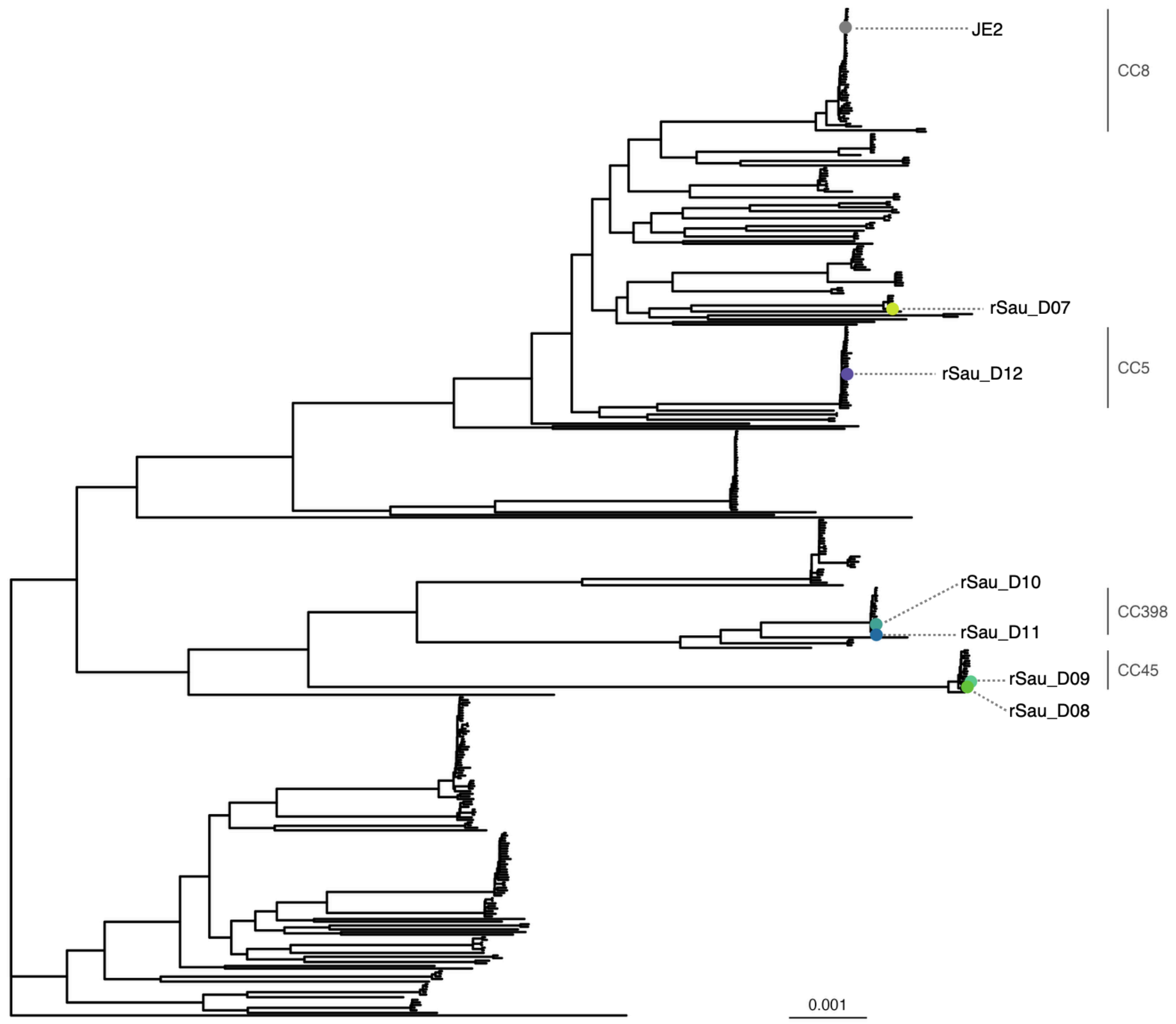


**Supplementary Figure 2: Phylogenetic tree showing the genomes of resident *S. aureus* (rSau) from D07-D12 in the context of a curated set of 334 *S. aureus* genomes from the Staphopia database.** To investigate the phylogenetic relatedness of the resident *S. aureus* isolates in a wider context, we downloaded a curated set of 334 *S. aureus* genomes originating from the Staphopia database, called non-redundant diversity (NRD) set [2,3], where each genome represents a unique sequence type (ST). We used the core gene alignment from a pangenome including our resident *S. aureus* isolates, our focal strain MRSA USA300 JE2, and the NRD data set obtained with Roary v3.13.0 [4] to construct a maximum-likelihood tree using IQ-TREE v3.0.1 [5]. We determined the sequence type with mlst v2.23.0

(https://github.com/tseemann/mlst) incorporating components of the pubMLST website (https://pubmlst.org/) [6] and assigned the sequence types to the corresponding clonal complexes (CC) based on pubMLST. Five out of six resident *S. aureus* genomes were assigned to known clonal complexes (grey labels on the right). More information about these strains is provided in Supplementary Table 1, and details on the whole-genome sequencing are described in Supplementary Methods.

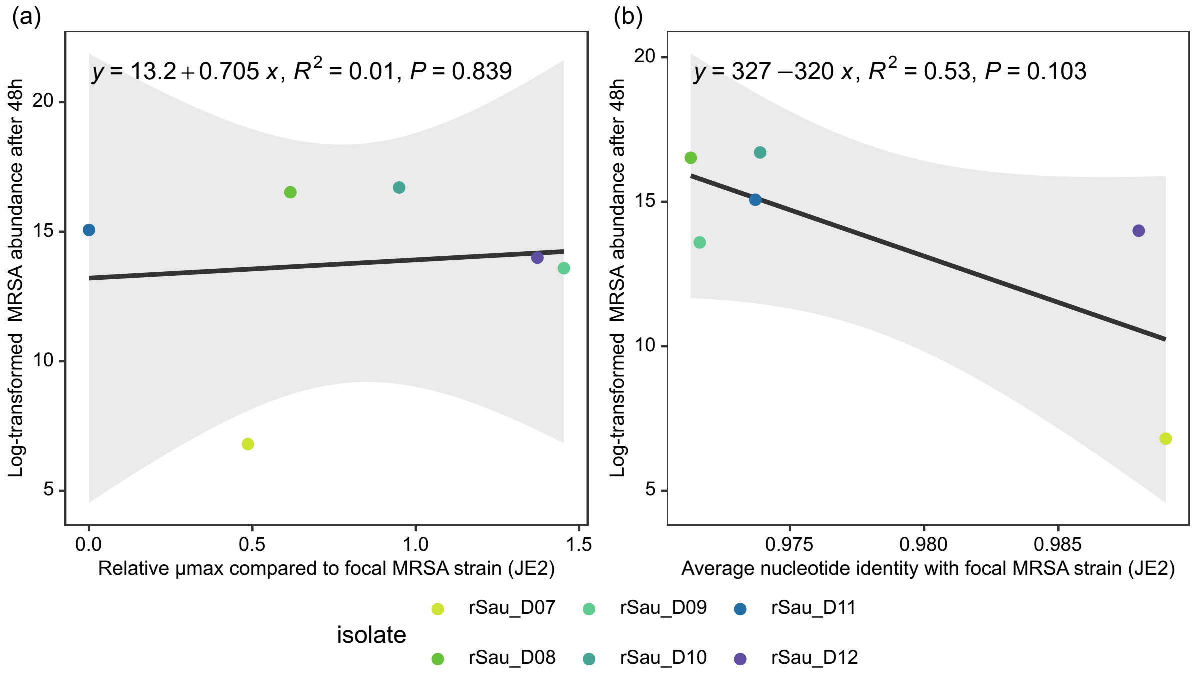


**Supplementary Figure 3: (a) Linear regression of MRSA inhibition by resident *S. aureus* isolates and their relative maximum growth rate ($\mu$max) compared to focal MRSA strain JE2.** To monitor growth of resident *S. aureus* isolates and MRSA strain JE2 in pure culture in basal medium, we used a plate reader (Tecan NanoQuant Infinite M200 Pro), shaking and measuring optical density at 600 nm every 10 min during 48 h. We used the fit_easylinear function from growthrates package v0.8.5 in R [7] to fit an exponential growth model with a time window of 10 h and extracted maximum growth rates. Individual points show the median relative maximum growth rates compared with focal MRSA strain JE2 from three replicates. **(b) Linear regression of MRSA inhibition by resident *S. aureus* isolates and their average nucleotide identity (ANI) with focal MRSA strain JE2.** We calculated ANI between the genome of each resident *S. aureus* isolate and the genome of the focal MRSA strain JE2 using pyani [8]. We found no significant correlation between the ANI to the focal MRSA and log-transformed median MRSA abundance in pairwise co-culture with each resident *S. aureus* isolate (see Fig. 3a) with a linear regression model ($p$ = 0.103), although the isolate with the highest ANI to the focal strain also caused the strongest inhibition (rSau_D07). Overall, we detected no significant phylogenetic structure for the degree of MRSA inhibition by different isolates (Pagel's $\lambda = 0.40$, $p = 0.67$). Note that statistical power of these tests are limited by sample size ($n = 6$).

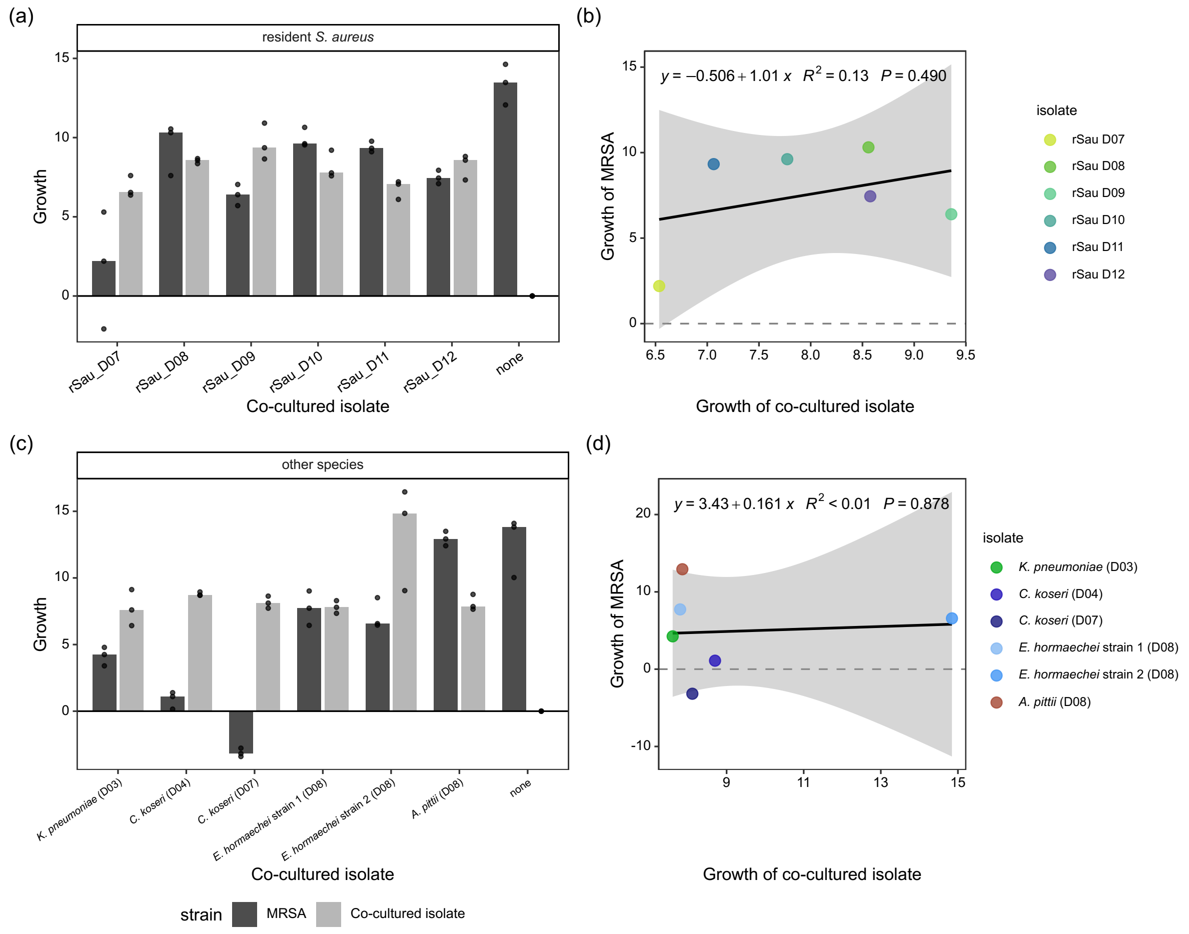


**Supplementary Figure 4: Net population growth of MRSA and resident *S. aureus* isolates and Enterobacteriaceae isolates in pairwise co-culture.** We calculated the Malthusian growth parameter ($\ln(\frac{\text{final abundance}}{\text{inoculum}})$) [1] for both the focal MRSA and the co-cultured isolate using initial and final abundances assessed by plating the same dilutions on both MRSA- and (a, b) *S. aureus*-selective chromatic agar, calculating resident *S. aureus* abundance by subtracting MRSA abundance from total *S. aureus* counts, or (c, d) Muller-Hinton chromatic agar. In (a) and (c), the individual replicates are represented by points, bars indicate the median (n = 3). The linear regressions in (b) and (d) are based on the median across the three replicates.

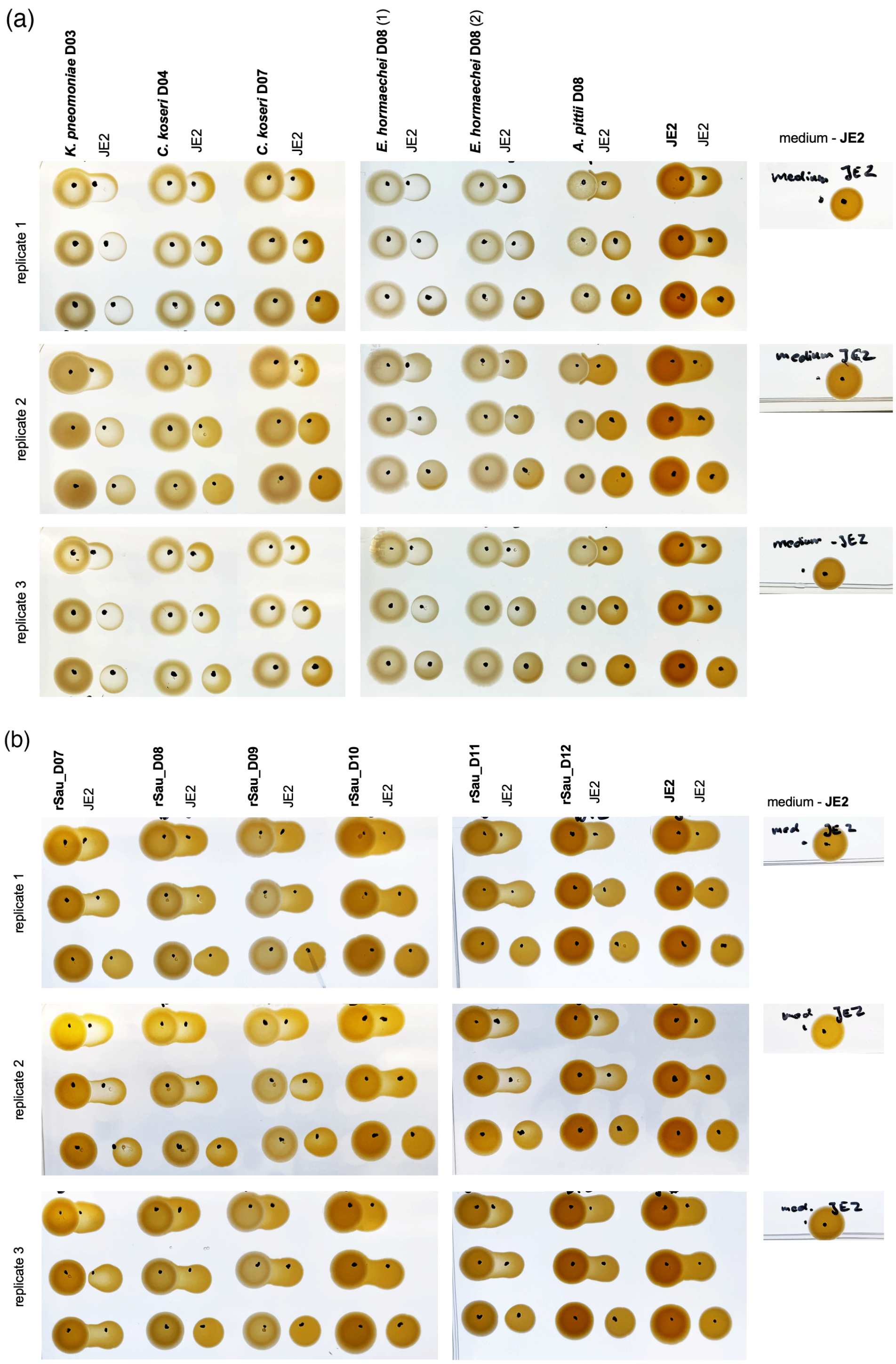


**Supplementary Figure 5: Pairwise inhibition in a side-by-side co-culture assay on BHI agar.** We pre-grew spots of 100-fold diluted overnight cultures of (a) Enterobacteriaceae isolates and the non-Enterobacteriaceae isolate *Acinetobacter pittii* (for reference), or (b) resident *S. aureus*, overnight on BHI agar. The next day, we added 100-fold diluted MRSA overnight culture with increasing difference next to the pre-grown isolate spots. Each assay was performed in 3 biological replicates.

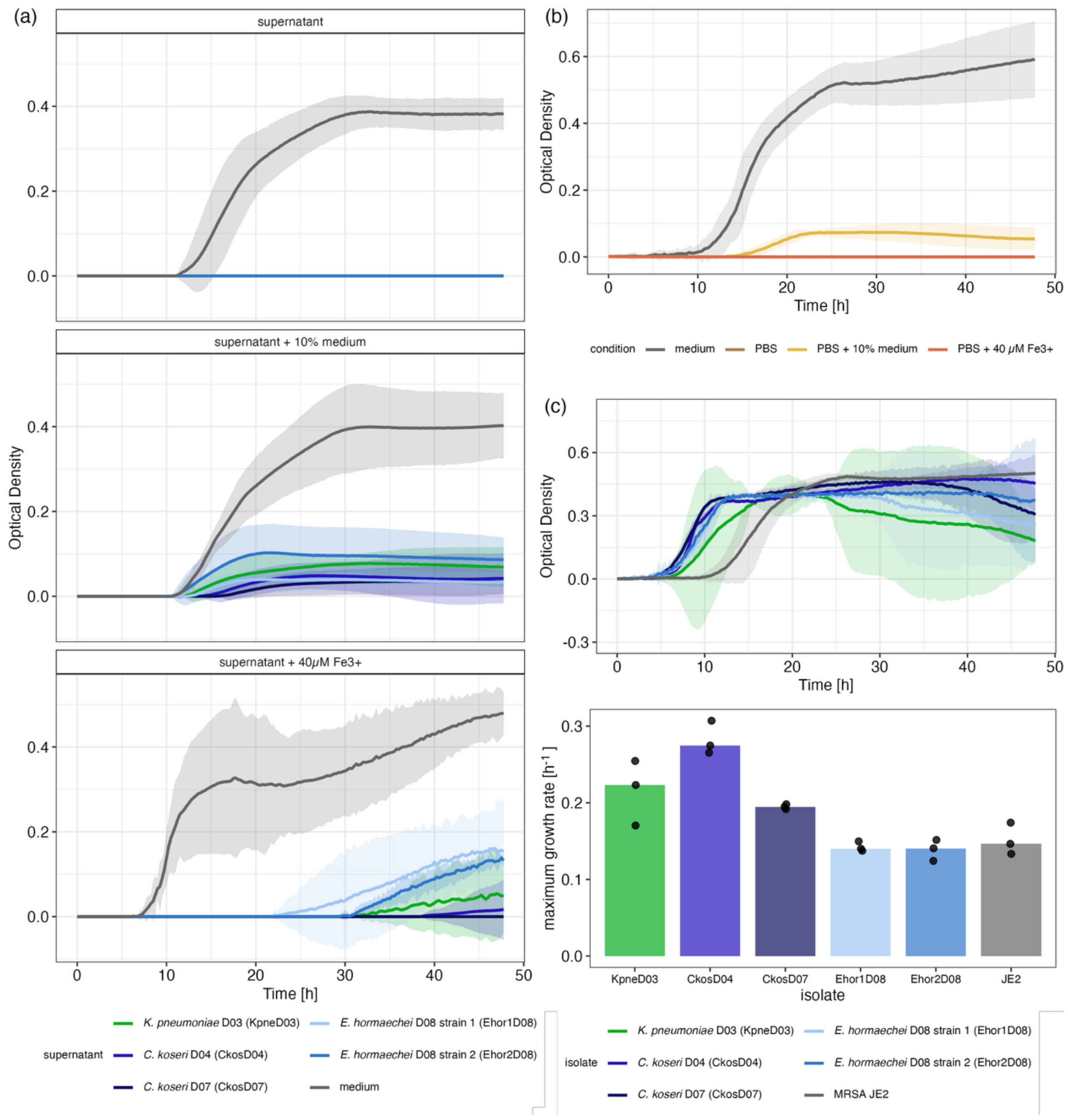


**Supplementary Figure 6: (a) Growth of MRSA in Enterobacteriaceae supernatants with varying nutrient availability.** We obtained supernatants by sterile-filtering cultures of Enterobacteriaceae isolates in basal medium after 48 h. The three panels show focal MRSA strain JE2 growth in unmodified supernatants (top), supernatants supplemented with 10% fresh basal medium (middle) or with 40 $\mu$M $Fe^{3+}$ (bottom). MRSA culture in fresh basal medium instead of culture supernatants, supplemented accordingly, is shown as reference (grey). Focal MRSA growth in unmodified Enterobacteriaceae supernatants was fully inhibited, but partially restored by addition of 10% fresh basal medium. When the supernatants were supplemented with 40 $\mu$M $Fe^{3+}$, a known limited resource in the nose underlying previously characterised competitive interactions among microbiota [9–12], did not restore MRSA growth in Enterobacteriaceae supernatants (Fig. S6a). MRSA remained unable to initiate exponential growth for more than 20 h, and the eventual increase in biomass occurred at similar levels in the control and iron-supplemented supernatants. **(b) Replacing supernatants with buffer results in similar MRSA growth.** In a follow-up experiment, we replaced Enterobacteriaceae supernatants with buffer (PBS). Growth of MRSA was not detectable in PBS alone (brown) but was partially restored when PBS was supplemented with 10% basal medium (yellow). Thus, supplementation with nutrients had similar effects on MRSA growth in both Enterobacteriaceae supernatants and PBS, suggesting low nutrient levels may explain MRSA inhibition in (a). **(c) Growth of Enterobacteriaceae isolates in pure culture.** Top panel shows growth curves of Enterobacteriaceae isolates and focal MRSA strain JE2 (as reference) in basal medium, and bottom panel shows corresponding maximum growth rates. We used the same inocula as in Fig. 3b, meaning that MRSA was inoculated at 100-fold lower starting density compared to the Enterobacteriaceae isolates. This explains the longer lag-phase of MRSA. To obtain growth curves, we used a plate reader (Tecan NanoQuant Infinite M200 Pro), shaking and measuring optical density at 600 nm every 10 min during 48 h. Growth curves are plotted as mean optical density (after background correction by subtracting the mean optical density of medium-only wells at each time point) with a 95%-confidence interval ($n = 3$). We used the fit_easylinear function from growthrates package v0.8.5 in R [7] to fit an exponential growth model with a time window of 10 h. We extracted maximum growth rates and normalised them by subtracting the maximum growth rate of medium-only wells. Bars on maximum growth rate plot show the median, points show individual replicates.

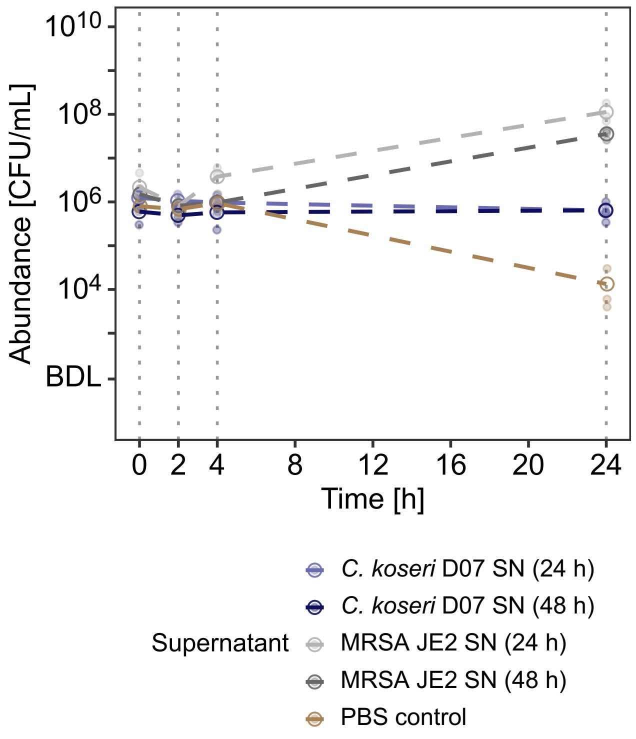


**Supplementary Figure 7: Sterile *C. koseri* D07 culture supernatant does not induce negative MRSA population growth.** We inoculated $10^6$ CFU/mL of MRSA (corresponding to the same inoculum used for Fig. 4) in sterile-filtered supernatants of *C. koseri* D07 cultures in basal medium, harvested after 24 h and 48 h. As comparison, we used MRSA culture supernatants and phosphate-buffered saline (labelled 'PBS control' in the legend). We measured MRSA abundances by plating serial dilutions directly after MRSA inoculation, and after 2, 4 and 24 h of incubation. Unlike our other supernatant assays using optical density to track biomass over time (Fig. S6), this would allow us to detect a decline in the bacterial abundance in CFU/mL over time. Filled points represent invidiual replicates ($n$ = 3), empty circles and dashed lines show median abundance over time.

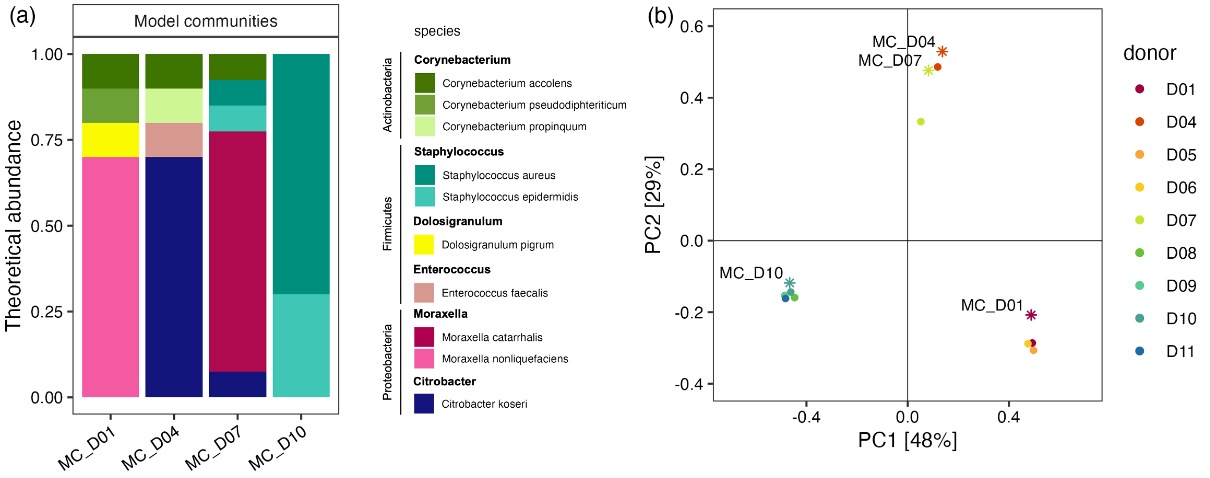


**Supplementary Figure 8: (a) Theoretical composition of model communities containing all bacterial isolates obtained from donors D01, D04, D07 or D10.** The theoretical relative abundance of the dominating species is 70%, and the remaining isolates make up a total of 30%. **(b) Principal coordinate analysis (PCoA) of the community composition.** We included the initial nasal swab samples (points, $n = 9$, Fig. 2a) and the theoretical composition of the model communities (stars, $n = 4$). The PCoA is based on species-level Bray-Curtis dissimilarity. Note that this does not account for taxonomic relatedness among species, but considers each taxon as completely independent. As a result, the distance MC_D07 and MC_D01 is relatively large, although they are both dominated by a (different) *Moraxella* species.

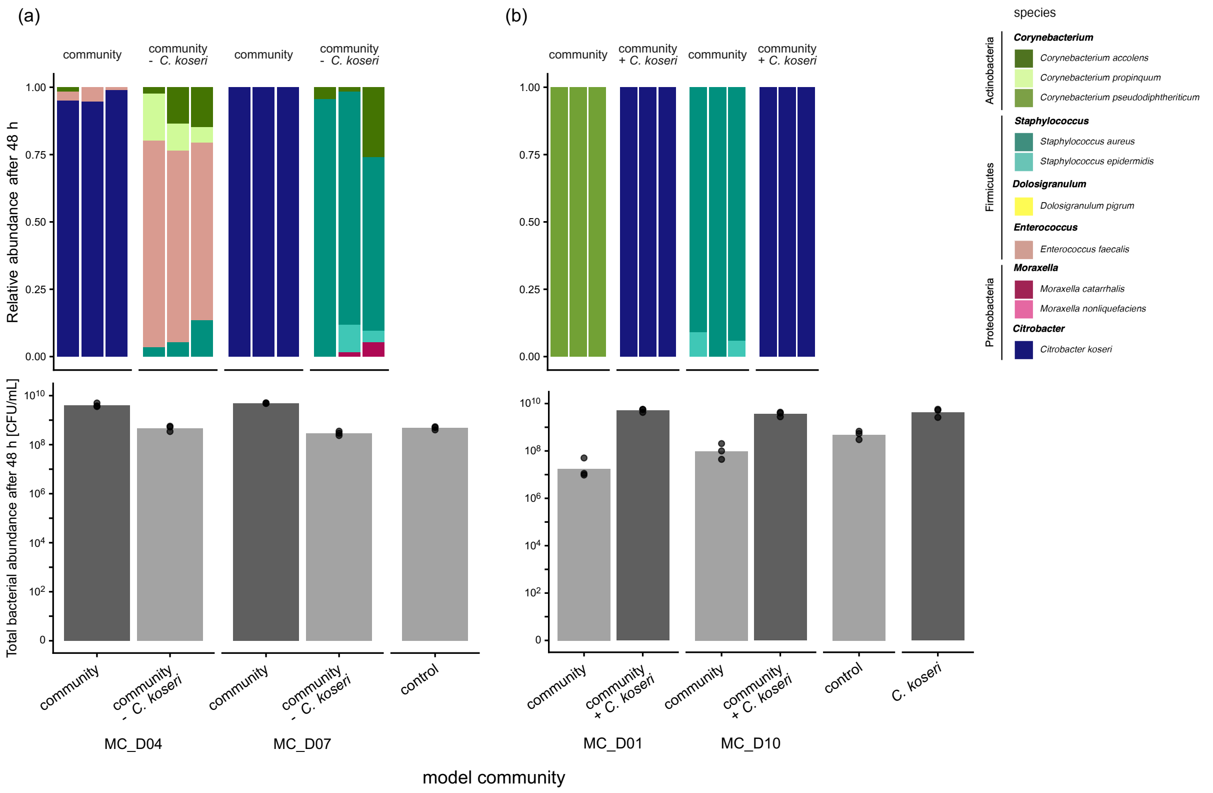


**Supplementary Figure 9: Final composition and total bacterial abundance in (a) model communities containing a resident *C. koseri* isolate, and a manipulated 'drop-out' version without *C. koseri,* and (b) model communities representing resident *C. koseri*-free samples, with 'drop-in' versions inoculated with the *C. koseri* isolate from D07.** Top panels show relative abundances of each species, bottom panels show total final bacterial abundance. Overall, presence of *C. koseri* was associated with higher final total bacterial abundance (two-way ANOVA on rank-transformed data, effect of *C. koseri* presence: $F_{2,10} = 50.3$, $p < 0.001$). All data shown here are based on colony counts on COS agar plates. Species differentiation is based on colony morphology. Note this may result in a detection bias towards more abundant species. Bars show medians, colour indicates presence (dark) or absence (bright) of *C. koseri*, points show individual microcosms ($n = 3$).

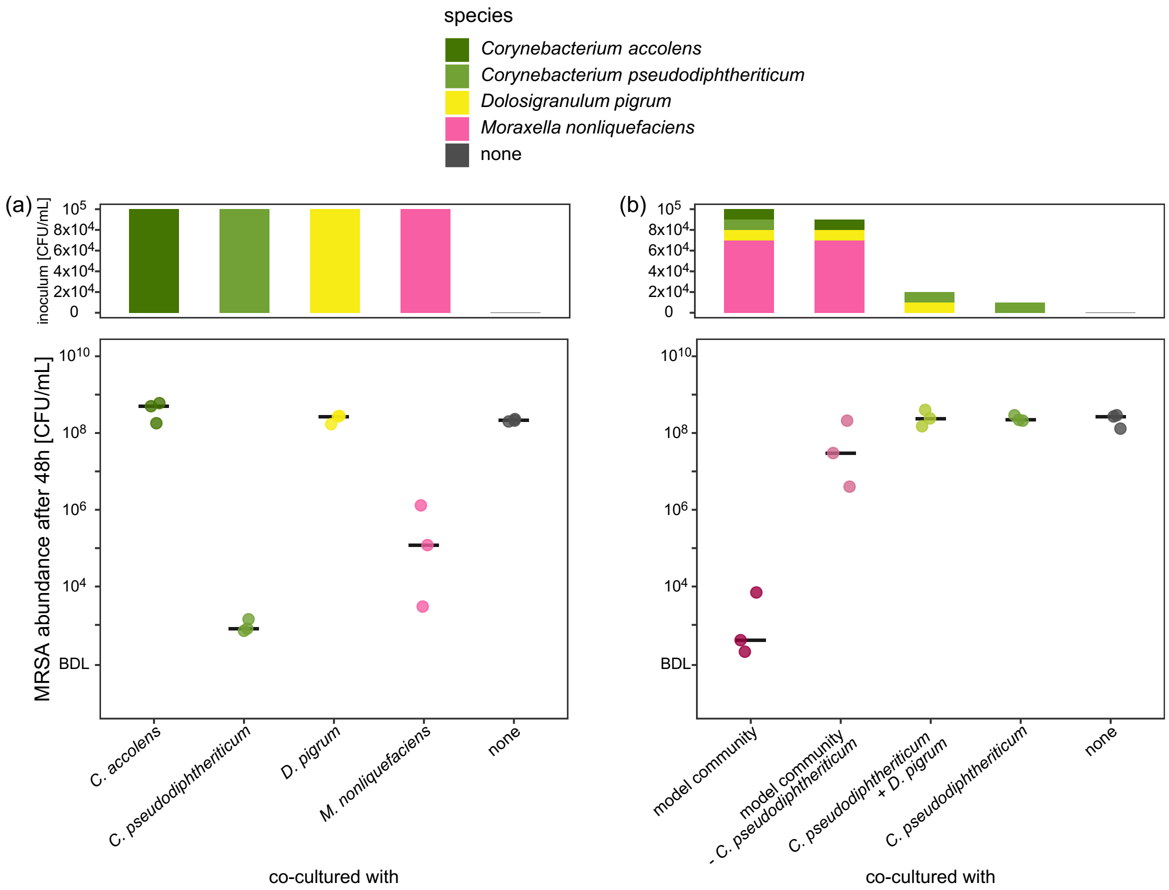


**Supplementary Figure 10: *C. pseudodiphtheriticum* is the major driver of colonisation resistance to MRSA in MC_D01.** Initial community composition is displayed in the top panels (showing the target abundance of each species upon inoculation; see Methods), and final MRSA abundance in the microcosms is shown in the bottom panels. **(a) Individual bacterial isolates from D01 exhibited varying MRSA inhibition in pairwise co-culture.** Final MRSA abundance varied among co-culture treatments with each isolate inoculated at a starting density of $10^5$ CFU/mL (one-way ANOVA on rank-transformed data: $F_{4,10} = 10.15$, $p < 0.01$), being lowest with *C. pseudodiphtheriticum* and also reduced with *M. nonliquefaciens*. **(b) *C. pseudodiphtheriticum* is essential for MRSA inhibition by MC_D01.** As in Fig. 5, the complete MC_D01 potently inhibits MRSA growth. Removal of *C. pseudodiphtheriticum* increased final MRSA abundance $7.5\times10^4$-fold compared to the complete community (unpaired two-sample *t*-test on log-transformed final MRSA abundance after correction for multiple testing by sequential Bonferroni: $p < 0.001$), reaching levels not significantly different from those in pure culture (unpaired two-sample $t$-test on log-transformed final MRSA abundance after correction for multiple testing by sequential Bonferroni: $p > 0.05$). When inoculated at the same density as in the model community ($10^4$ CFU/mL) as opposed to the higher inoculum size in (a), *C. pseudodiphtheriticum* alone, or in combination with *D. pigrum,* did not reduce final MRSA abundance compared to pure culture.

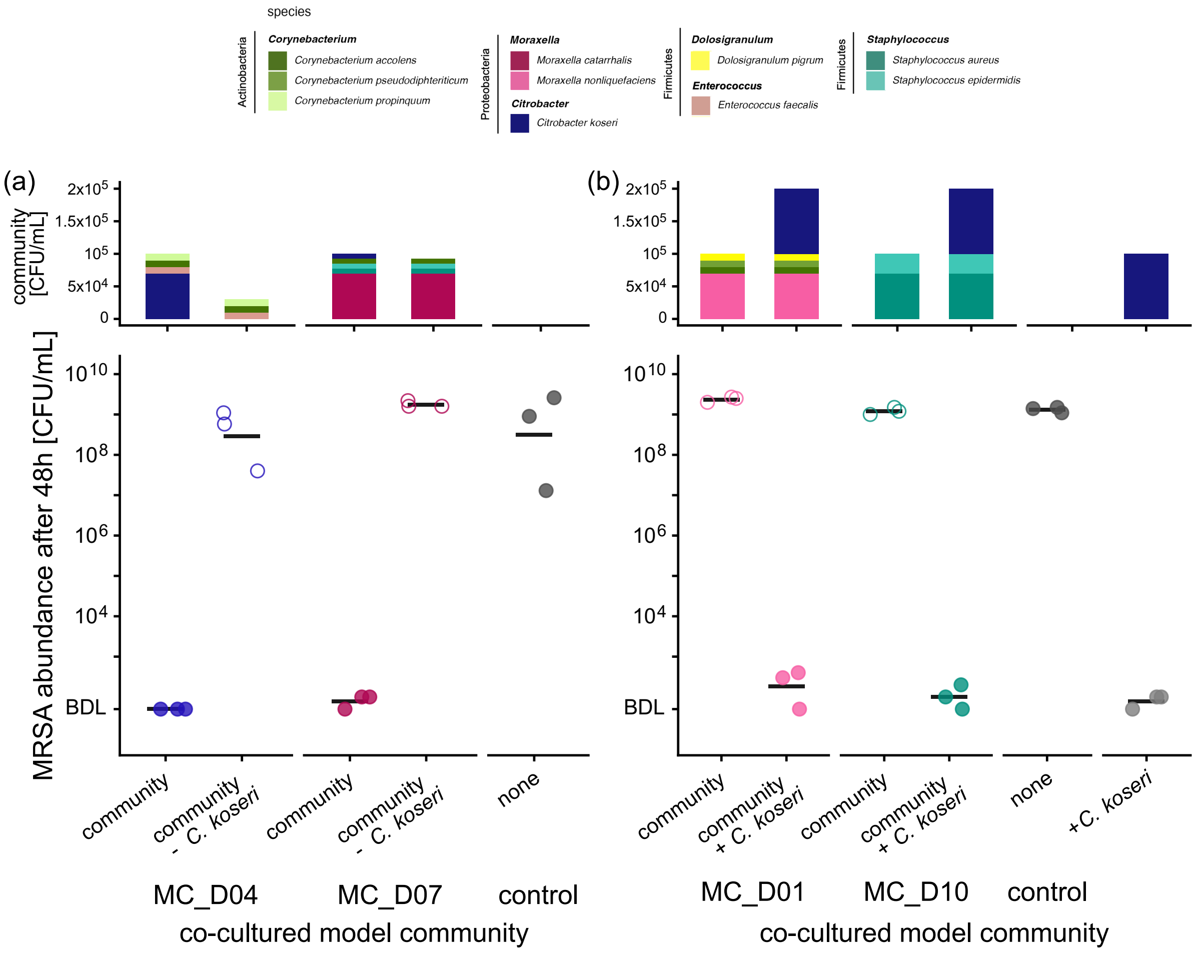


**Supplementary Figure 11: *C. koseri* drives community-level colonisation resistance to MRSA of nasal model communities in synthetic nasal medium.** To test the robustness of our previous findings using basal medium (RPMI + 1% BHI; Fig. 5), we quantified the effect of (a) removing resident *C. koseri* and (b) introducing foreign *C. koseri* on community-level colonisation resistance to MRSA in synthetic nasal medium (SNM3, prepared as described previously [13]). As in basal medium, *C. koseri* was the key driver of colonisation resistance: in (a), removal of resident *C. koseri* increased MRSA abundance (two-way ANOVA on rank-transformed data, effect of *C. koseri* presence: $F_{1,9}$ = 94.8, $p$ < 0.001); in (b), addition of *C. koseri* reduced MRSA abundance (two-way ANOVA on log-transformed data, effect of *C. koseri* presence: $F_{1,9}$ = 1985.8, $p$ < 0.001). In absence of *C. koseri,* all communities supported high MRSA growth in SNM3. This contrasts with the high baseline colonisation resistance of

MC_D01 in basal medium, and the intermediate colonisation resistance of MC_D10 (without addition of *C. koseri*; Fig. 5), showing some assembled communities were more inhibitory in basal medium, although the effect of *C. koseri* was consistent.

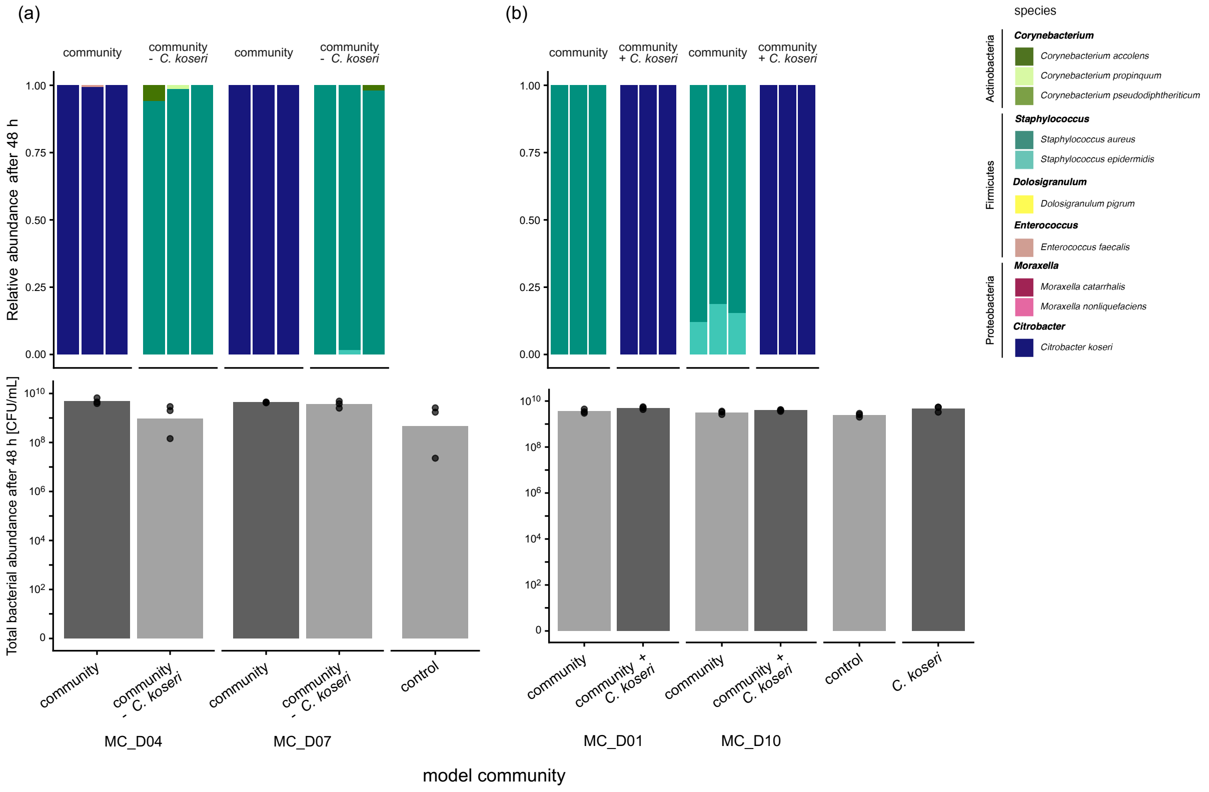


**Supplementary Figure 12: Final composition and total bacterial abundance in model communities in synthetic nasal medium.** (a) Model communities containing a resident *C. koseri* isolate, and a manipulated 'drop-out' version without *C. koseri,* and (b) model communities representing resident *C. koseri*-free samples, with 'drop-in' versions inoculated with the *C. koseri* isolate from D07. Top panels show relative abundances of each species, bottom panels show total final bacterial abundance (after 48 h of incubation). All data shown here are based on colony counts on COS agar plates. Species differentiation is based on colony morphology. Note this may result in a detection bias towards more abundant species. Bars show medians, colour indicates presence (dark) or absence (bright) of *C. koseri*, points show individual microcosms ($n$ = 3). Initial community composition and MRSA abundance after 48 h of the same microcosms are shown in Fig. S11.

### Supplementary Tables

| Isolate | *agr* type | MLST | Clonal complex |
|---|---|---|---|
| rSau_D07 | 1 | 3035 | unassigned |
| rSau_D08 | 1 | 46 | CC45 |
| rSau_D09 | 1 | 508 | CC45 |
| rSau_D10 | 1 | 398 | CC398 |
| rSau_D11 | 1 | 398 | CC398 |
| rSau_D12 | 2 | 5 | CC5 |
| JE2 | 1 | 8 | CC8 |

**Supplementary Table 1: Characteristics of resident *S. aureus* isolates from donors D07-D12 based on whole-genome sequencing**. We used the following bioinformatic tools: AgrVATE for *agr*-type [3], mlst for MLST (Seemann T, mlst Github https://github.com/tseemann/mlst, unpublished) incorporating components of the pubMLST website (https://pubmlst.org/) [6], and assigned the sequence types to the corresponding clonal complexes based on pubMLST.

| Model community | Origin | Native resident Ckos | Condition | Species | Pre-inoculum type | Pre-inoculum abundance [CFU / mL] | Target abundance in MC [CFU / mL] |
|---|---|---|---|---|---|---|---|
| MC_D01 | D01 | no | community | ***Moraxella nonliquefaciens*** | suspension | 1.50E+07 | 7.00E+04 |
| MC_D01 | D01 | no | community | *Corynebacterium accolens* | suspension | 1.07E+07 | 1.00E+04 |
| MC_D01 | D01 | no | community | *Corynebacterium pseudodiphtheriticum* | suspension | 1.37E+07 | 1.00E+04 |
| MC_D01 | D01 | no | community | *Dolosigranulum pigrum* | suspension | 8.67E+05 | 1.00E+04 |
| MC_D01 | D01 | no | community + *C. koseri* | ***Moraxella nonliquefaciens*** | suspension | 1.50E+07 | 7.00E+04 |
| MC_D01 | D01 | no | community + *C. koseri* | *Corynebacterium accolens* | suspension | 1.07E+07 | 1.00E+04 |
| MC_D01 | D01 | no | community + *C. koseri* | *Corynebacterium pseudodiphtheriticum* | suspension | 1.37E+07 | 1.00E+04 |
| MC_D01 | D01 | no | community + *C. koseri* | *Dolosigranulum pigrum* | suspension | 8.67E+05 | 1.00E+04 |
| MC_D01 | D07 | no | community + *C. koseri* | *Citrobacter koseri* (drop-in) | liquid culture | 5.33E+09 | 1.00E+05 |
| MC_D04 | D04 | yes | community | ***Citrobacter koseri*** | liquid culture | 8.67E+09 | 7.00E+04 |
| MC_D04 | D04 | yes | community | *Enterococcus faecalis* | liquid culture | 6.33E+08 | 1.00E+04 |
| MC_D04 | D04 | yes | community | *Corynebacterium accolens* | suspension | 1.87E+07 | 1.00E+04 |
| MC_D04 | D04 | yes | community | *Corynebacterium propinquum* | suspension | 1.90E+07 | 1.00E+04 |
| MC_D04 | D04 | yes | community - *C. koseri* | *Enterococcus faecalis* | liquid culture | 6.33E+08 | 1.00E+04 |
| MC_D04 | D04 | yes | community - *C. koseri* | *Corynebacterium accolens* | suspension | 1.87E+07 | 1.00E+04 |
| MC_D04 | D04 | yes | community - *C. koseri* | *Corynebacterium propinquum* | suspension | 1.90E+07 | 1.00E+04 |
| MC_D07 | D07 | yes | community | ***Moraxella catarrhalis*** | suspension | 7.33E+06 | 7.00E+04 |
| MC_D07 | D07 | yes | community | *Corynebacterium accolens* | suspension | 1.87E+07 | 7.50E+03 |
| MC_D07 | D07 | yes | community | *Staphylococcus aureus* | liquid culture | 4.33E+09 | 7.50E+03 |
| MC_D07 | D07 | yes | community | *Staphylococcus epidermidis* | liquid culture | 1.83E+09 | 7.50E+03 |
| MC_D07 | D07 | yes | community | *Citrobacter koseri* | liquid culture | 5.33E+09 | 7.50E+03 |
| MC_D07 | D07 | yes | community - *C. koseri* | ***Moraxella catarrhalis*** | suspension | 7.33E+06 | 7.00E+04 |
| MC_D07 | D07 | yes | community - *C. koseri* | *Corynebacterium accolens* | suspension | 1.87E+07 | 7.50E+03 |
| MC_D07 | D07 | yes | community - *C. koseri* | *Staphylococcus aureus* | liquid culture | 4.33E+09 | 7.50E+03 |
| MC_D07 | D07 | yes | community - *C. koseri* | *Staphylococcus epidermidis* | liquid culture | 1.83E+09 | 7.50E+03 |
| MC_D10 | D10 | no | community | ***Staphylococcus aureus*** | liquid culture | 1.40E+09 | 7.00E+04 |
| MC_D10 | D10 | no | community | *Staphylococcus epidermidis* | liquid culture | 2.61E+08 | 3.00E+04 |
| MC_D10 | D10 | no | community + *C. koseri* | ***Staphylococcus aureus*** | liquid culture | 1.40E+09 | 7.00E+04 |
| MC_D10 | D10 | no | community + *C. koseri* | *Staphylococcus epidermidis* | liquid culture | 2.61E+08 | 3.00E+04 |
| MC_D10 | D07 | no | community + *C. koseri* | *Citrobacter koseri* (drop-in) | liquid culture | 5.33E+09 | 1.00E+05 |

**Supplementary Table 2: Composition of model communities (visualised in Fig. 5).** Each model community (MC) consists of multiple species originating from the same nasal swab sample. To assemble each model community inoculum, we pre-cultivated the constituent isolates in pure liquid cultures in basal medium for 24 h. We found some isolates did not grow sufficiently in pure liquid cultures but did grow successfully on agar. For these isolates, instead of pre-cultivation in liquid culture we picked colonies from agar plates (incubated for 36 h), suspended them in basal medium and normalised to $OD_{600nm}$ = 1.0. Using defined isolate-specific dilution factors based on previously assessed bacterial densities of all pre-inocula, we assembled 10X concentrated model community inocula in basal medium. Total target abundance of each unmanipulated model community (condition = "community") was $10^5$ CFU/mL in nasal microcosms, with the dominating species (bold) at a fixed relative abundance of 70% and the other isolates making up the remaining 30%. Model communities lacking a native resident *C. koseri* were inoculated with $5\times10^5$ CFU/mL of the *C. koseri* isolate from D07 (blue; condition = "community + *C. koseri*). We inoculated microcosms containing basal medium and Amies Transport Medium (final ratio 4:1) with each model community (with / without *C. koseri*, $n = 3$ per condition and community). After 2 h of pre-conditioning at 37° C, we added 10 $\mu$L of 1:10'000 diluted overnight culture of MRSA strain JE2 in PBS (separate cultures for each replicated microcosm) and aliquoted 130 $\mu$L for plating of timepoint 0 h on COS and MRSA selective agar, resulting in a final volume of 1 mL per microcosm. We incubated all microcosms at 37° C for 48 h and then plated serial dilutions of each microcosm on MRSA selective agar and 50 $\mu$L on COS plates using glass beads.

| gene | D03Kpne | D04Ckos | D07Ckos | D08Ehor1 | D08Ehor2 |
|---|---|---|---|---|---|
| T6SSi_evpJ | 0 | 0 | 3 | 4 | 4 |
| T6SSi_tssA | 2 | 0 | 1 | 2 | 2 |
| T6SSi_tssB | 1 | 0 | 1 | 2 | 2 |
| T6SSi_tssC | 0 | 0 | 0 | 0 | 0 |
| T6SSi_tssD | 1 | 0 | 8 | 5 | 5 |
| T6SSi_tssE | 1 | 0 | 1 | 2 | 2 |
| T6SSi_tssF | 1 | 0 | 1 | 2 | 2 |
| T6SSi_tssG | 1 | 0 | 1 | 2 | 2 |
| T6SSi_tssH | 2 | 0 | 2 | 3 | 4 |
| T6SSi_tssI | 1 | 0 | 2 | 3 | 3 |
| T6SSi_tssJ | 1 | 0 | 1 | 2 | 2 |
| T6SSi_tssK | 1 | 0 | 1 | 2 | 2 |
| T6SSi_tssL | 1 | 0 | 1 | 2 | 2 |
| T6SSi_tssM | 1 | 0 | 1 | 2 | 2 |
| system wholeness | 0.857 | 0 | 0.929 | 0.929 | 0.929 |

**Supplementary Table 3: Type VI secretion system (T6SS$^{i}$) gene content across Enterobacteriaceae isolates.** T6SS$^{i}$ mediates contact-dependent killing of bacterial competitors [14,15]. We used TXSScan v1.1.3 [16] in MacSyFinder v2.1.5 [17,18] to detect mandatory T6SS$^{i}$ genes in short-read assembled Enterobacteriaceae isolate genomes (unordered replicon mode). Values represent number of hits for detected mandatory T6SS$^{i}$ genes and system wholeness indicates the proportion of mandatory genes with at least one hit for each isolate. Abbreviations: *K. pneumoniae* D03 (D03Kpne), *C. koseri* D04 / D07 (D04Ckos / D07Ckos), *E. hormaechei* D08 strain 1 / strain 2 (D08Ehor1 / D08Ehor2).

### Supplementary Methods

#### Whole-genome sequencing

Extraction and whole-genome sequencing of all resident *S. aureus* (short- and long-read sequencing) and Enterobacteriaceae (short-read sequencing) isolates was performed at the Institute of Medical Microbiology at the University of Zurich (IMM, UZH). Genomic DNA was isolated using the Maxwell RSC Blood DNA Kit (Promega, Madison, USA). For short-read sequencing, libraries were prepared with the QIAseq FX DNA Library Kit (QIAGEN, Hilden, Germany) and sequenced as 150 bp paired-end reads on a NextSeq 1000 system (Illumina). For additional long-read sequencing (only for resident *S. aureus* isolates), the Rapid Barcoding Kit V13 (Oxford Nanopore Technologies) was used for library preparation, sequencing was performed with the GridION instrument (Oxford Nanopore Technologies, MinION R9.3 flow cell). We inspected the quality of the sequences with FastQC v0.12.1 [19] and generated hybrid (for resident *S. aureus* isolates) or short-read (for Enterobacteriaceae isolates) assemblies with Unicycler [20]. We annotated the genome assemblies with Prokka v1.14.5 [21].
